# HIV Infection impairs the Host Response to *Mycobacterium tuberculosis* Infection by altering Surfactant Protein D function in the Human Lung Alveolar Mucosa

**DOI:** 10.1101/2023.10.01.560171

**Authors:** Anwari Akhter, Juan I. Moliva, Abul K. Azad, Angélica Olmo-Fontánez, Andreu Garcia-Vilanova, Julia M. Scordo, Mikhail A. Gavrilin, Phillip T. Diaz, Janice J. Endsley, Susan T. Weintraub, Larry S. Schlesinger, Mark D. Wewers, Jordi B. Torrelles

## Abstract

Tuberculosis is the leading cause of death for people living with HIV (PLWH). We hypothesized that altered functions of innate immune components in the human alveolar lining fluid of PLWH (HIV-ALF), drive susceptibility to *Mycobacterium tuberculosis* (*M.tb*) infection. Our results indicate a significant increase in oxidation of innate proteins and chemokine levels, and significantly lower levels and function of complement components and Th1/Th2/Th17 cytokines in HIV-ALF *vs.* control-ALF (non-HIV infected people). We further found a deficiency of surfactant protein-D (SP-D) and reduced binding of SP-D to *M.tb* that had been exposed to HIV-ALF. Primary human macrophages infected with *M.tb* exposed to HIV-ALF were significantly less capable of controlling the infection, which was reversed by SP-D replenishment in HIV-ALF. Thus, our data suggest that PLWH without antiretroviral therapy (ART) have declining host innate defense function in their lung mucosa, thereby favoring *M.tb* and potentially other pulmonary infections.

## INTRODUCTION

Worldwide, 38.4 million people are living with HIV/AIDS, including approximately 1.5 million people becoming newly infected with HIV annually, with a mortality rate of 650,000 people per year (1). Currently in the US, 50% of the people living with HIV (PLWH) are 50 years old or older (2); this number is predicted to be 70% by 2030 (3), indicating an increasingly high burden of HIV-infected individuals as people live longer with chronic HIV infection (4–9). Aging-related concerns in PLWH >50 years of age include accelerated CD4^+^ T cell loss, decreased immune recovery and increased risk of serious non-AIDS illnesses, further complicating antiretroviral therapy (ART) management.

The spectrum of lung diseases associated with HIV includes both infectious and non-infectious etiologies, but the mechanisms behind this are not well understood (10, 11). HIV infection results in immune dysregulation, dysfunction, and deficiency (12–14). These immunologic abnormalities are most marked in PLWH not receiving ART. Even though immune functions can partially restored in many PLWH receiving ART; persistent systemic inflammation driving their immunodeficiency may occur, particularly in PLWH who have lower CD4+ T cell counts at the time of ART initiation (15–18). Indeed, the chronic phase of HIV infection is associated with chronic immune activation and cell exhaustion, where recurrent episodes of respiratory infections (bacterial, viral, and fungal pneumonia) further increase chronic pulmonary inflammation and predispose PLWH to lung cancer (19).

Our understanding of the role of the lung alveolar environment in the development of these lung diseases in the current ART era is still limited. Oxidative stress levels in the healthy lung alveolar space are highly controlled throughout the majority of an individual’s life, typically increasing only with old age (20, 21) and progression of disease. At the initiation of the current study, we hypothesized that basal lung alveolar oxidation levels are increased in PLWH, driving immune dysfunction within the alveolar space. To test our hypothesis, we assessed lung alveolar space immune responses that are linked to oxidation and inflammation in HIV-negative control subjects and PLWH without ART. Our results indicate that PLWH have immune dysfunction in their alveolar lining fluid (ALF). This dysfunction is characterized by an increase in lung protein oxidation and chemokine levels, reduction of Th1/Th2/Th17 cytokines and growth factors in ALF. Furthermore, there was a reduction in complement and surfactant proteins A and D (SP-A and SP-D) levels and activity. The latter innate immune components play important roles in host responses to respiratory pathogens such as *Mycobacterium tuberculosis* (*M.tb*) (22–25). This innate immunodeficiency in the lungs of PLWH without ART was further validated by the observation that macrophages from healthy human donors were infected with *M.tb* that had been exposed to ALF from PLWH (HIV-ALF), there was an increase in both *M.tb* association with macrophages and in intracellular growth compared to *M.tb* exposed to control-ALF. This reduction in efficiency to control *M.tb* growth in macrophages was due to a decrease in phagosome maturation below baseline levels. This was reversed by addition of SP-D to HIV-ALF. Thus, our results indicate that innate dysfunction in the lung mucosa of PLWH could drive susceptibility to respiratory infections such as one caused by *M.tb*.

## MATERIALS AND METHODS

### Human subjects and ethics statement

All experimental procedures with human subjects were carried out in strict accordance with the US Code of Federal and Local Regulations, following The Ohio State University (OSU) and Texas Biomedical Research Institute (Texas Biomed) IRB approved protocols. Bronchoalveolar lavage fluid (BALF) to obtain ALF was collected from control and PLWH subjects. All subjects for this study provided informed written consent. We obtained BALF from age- and sex-matched healthy donors (median age 32, without comorbidities, non-smokers, non-drug users) and PLWH who were not receiving ART [undisclosed smoking, never or former smoking status, or drug use but currently declared non-smokers, median CD4^+^ T cell count of 378.5 cells/μl (93 – 661 cells/μl), with a median viral load of 54,812 RNA copies/ml (<100-210,504 RNA copies/ml), median age of 34.7 (23.2-50.9), and a median BMI of 18.82 (16.6-21.5), 90% male](20, 21, 26–30).

### Collection of human BALF and ALF

BALF was collected and ALF obtained at its physiological concentration present within the lung (at 1 – 1.5 mg of phospholipid/mL) as we previously described (21, 26–29, 31–33). Status of being either a never or a former smoker does not affect the phospholipid content in the donor’s lung mucosa (34). Concentrated ALF samples were stored in 50-to 100-μL aliquots at -80°C. Total protein content in ALF samples was determined by using a BCA protein assay kit (Thermo Fisher).

### Isolation and culture of human monocyte-derived macrophages

Peripheral blood mononuclear cells (PBMCs) were isolated from healthy donor blood and monocyte-derived macrophages (MDMs) were prepared from PBMCs as described (27, 31). Briefly, heparinized blood was layered on a Ficoll-Paque cushion (GE Healthcare, Uppsala, Sweden) to allow for the collection of PBMCs. PBMCs were then cultured in RPMI (Life Technologies, Carlsbad, CA) with 20% autologous serum in Teflon wells (Savillex, Eden Prairie, MN) for 5 days at 37°C/5% CO_2_ to allow for monocyte maturation into macrophages. PBMCs were harvested and MDMs purified by their adhesion to tissue culture dishes for 2 h in RPMI with 10% autologous serum, followed by washing away the lymphocytes. MDMs were then incubated overnight in RPMI with 10% autologous serum before their use in the downstream experiments.

### *M.tb* strains and culture

*M*.*tb* H_37_R_v_ (ATCC# 27294, Manassas, VA), GFP-*M.tb* Erdman (kindly provided by Dr. Marcus Horwitz, University of California, Los Angeles, CA) and *M.tb* Erdman-Red Cherry tomato (kindly provided by Dr. William Jacobs Jr., Albert Einstein Institute, New York, NY) were grown as we previously described (31). *M*.*tb* H_37_R_v_-Lux was kindly provided by Dr. Larry S. Schlesinger (Texas Biomed, San Antonio,TX) (35). SSB-GFP, *smyc’*::mCherry *M.tb* Erdman (kindly provided by Dr. David Russell, Cornell University, Ithaca, NY) was grown as described (32, 36). *M.tb* strains were cultured on 7H11 agar media with and without applicable antibiotics for 12-14 days at 37°C, 5% CO_2_.

### Determination of protein oxidation markers in human ALF

All ALF samples were normalized by protein content. Levels of protein carbonyls [a stable marker of reactive oxygen species (ROS)-induced oxidation] in ALF (using 8 μg of protein) was determined using OxiSelect Protein Carbonyl ELISA Kit (Cell Biolabs, Inc.). Levels of 3-nitrotyrosine-containing proteins [a stable marker of reactive nitrogen species [RNS]-induced oxidation) in ALF (using 10 μg of protein) were determined using OxiSelect Nitrotyrosine ELISA Kit (Cell Biolabs, Inc., San Diego, CA). Myeloperoxidase (MPO, indicative of tissue oxidation) levels in ALF (using 10 μg of protein) were determined by using LUMINEX (R&D Systems, Minneapolis, MN).

### Determination of immune Mmediators in human ALF

ALF samples were normalized by total protein content. Protein levels in ALFs were determined using custom Human Magnetic Luminex Assays (R&D Systems) for IFNγ, TNF, IL-12/23p40, IL-2, IL-4, IL-10, IL-6, IL-13, IL-17, IL-21, IL-22, IL-27, GM-CSF, TREM-1, IL-8, CCL2, complement component (C) 2, C5/C5a, C9, SP-D, and mannose binding lectin (MBL). SP-A levels were measured by Human SP-A ELISA kit (LifeSpan Biosciences, Inc., Seattle, WA) and C3 levels by Human Complement 3 ELISA kit (Abcam, Cambridge, MA) as per manufacturer instructions.

### Proteomic analyses of human ALF

ALF sample aliquots corresponding to 10 µg protein (EZQ™ Protein Quantitation Kit; Thermo Fisher) were mixed with 5% SDS/50 mM triethylammonium bicarbonate in the presence of protease and phosphatase inhibitors (Halt; Thermo Scientific). Then, these ALF samples were reduced with tris(2-carboxyethyl)phosphine hydrochloride, alkylated in the dark with iodoacetamide, and applied to S-Traps (micro; Protifi) for tryptic digestion (sequencing grade; Promega, Madison, WI) in 50 mM TEAB. Peptides were eluted from the S-Traps with 0.2% formic acid in 50% aqueous acetonitrile and quantified using Pierce™ Quantitative Fluorometric Peptide Assay (Thermo Scientific).

Data-independent acquisition mass spectrometry was conducted on an Orbitrap Fusion Lumos mass spectrometer (Thermo Scientific). On-line HPLC separation was accomplished with an RSLC NANO HPLC system (Thermo Scientific/Dionex): column, PicoFrit™ (New Objective; 75 μm i.d.) packed to 15 cm with C18 adsorbent (Vydac; 218MS 5 μm, 300 Å); mobile phase A, 0.5% acetic acid (HAc)/0.005% trifluoroacetic acid (TFA) in water; mobile phase B, 90% acetonitrile/0.5% HAc/0.005% TFA/9.5% water; gradient 3 to 42% B in 120 min; flow rate, 0.4 μL/min. A pool was made of all of the samples, and 1-µg peptide aliquots were analyzed using gas-phase fractionation and 4-*m/z* windows (30k resolution for precursor and product ion scans, all in the orbitrap) to create a DIA chromatogram library by searching against a panhuman spectral library (37, 38). Experimental samples were randomized for sample preparation and analysis. Injections of 1 µg of peptides were employed. MS data for experimental samples were acquired in the orbitrap using 12-*m/z* windows (staggered; 30k resolution for precursor and product ion scans) and searched against the chromatogram library. Scaffold DIA (v3.3.1 Proteome Software, Portland, OR) was used for all DIA data processing.

### Preparation of human ALF-exposed *M.tb*

*M.tb* exposure to human ALF was performed as described (26–29, 31, 33). Briefly, *M.tb* single cell suspensions were prepared and counted as previously described, and 1×10^8^ bacteria were exposed to physiological concentrations of ALF (50 μL as described above) for 12 h at 37°C, 5% CO_2_. After exposure, ALF-exposed *M.tb* bacilli were washed twice with sterile endotoxin-free saline and immediately used for analysis and *in vitro* infections (see below) (31, 39, 40).

### Determination of functionality of ALF innate proteins by *M.tb*-protein binding assays

Human ALF-exposed-*M.tb* was washed with TBS + 2 mM CaCl_2_ (TBS-C) to remove residual ALF and subsequently suspended in 500 μL of TBS-C. Human ALF-exposed *M.tb* (1×10^7^) was added to triplicate wells of 96-well plates and dried overnight. Binding of SP-A, SP-D, C3, and MBL contained in human ALF samples to *M.tb* was determined by indirect ELISA using primary monoclonal antibodies (mouse anti-human SP-A, mouse anti-human SP-D, mouse anti-human C3/C3b, and mouse anti-human MBL; all from Abcam) and secondary antibodies (donkey anti-mouse HRP and goat-anti rabbit HRP; Santa Cruz). Amounts of protein binding to *M.tb* were quantified by colorimetric measurements by absorbance at 450 nm using a GloMax plate reader (Version 3.0; Promega Corporation), and plotted based on absorbance values relative to the baseline (negative control, No ALF).

### ALF-exposed *M.tb* association with, and trafficking, replication and survival within human macrophage

To determine *M.tb-*macrophage association, GFP-*M.tb* Erdman and Red-Cherry tomato *M.tb* Erdman bacteria were exposed to HIV-ALF or control-ALF, respectively, washed to remove all traces of ALF before both bacterial strains were mixed at 1:1 ratio. MDM monolayers on coverslips were then infected with the bacterial mixture at an MOI of 10:1 for 2 h. Coverslips were fixed with 4% paraformaldehyde, followed by mounting with ProLong diamond antifade mount with DAPI (Invitrogen, Waltham, MA) (31, 33). Coverslips were imaged by using a Zeiss LSM 800 Confocal Microscope. At least 300 bacteria were quantified per condition. This study was repeated by switching the ALF exposure: GFP-*M.tb* Erdman was exposed to control-ALF and Red-Cherry tomato *M.tb* Erdman to HIV-ALF. In both cases we observed the same phenotype. To determine intracellular trafficking, MDM monolayers on coverslips were infected with GFP-*M.tb* Erdman exposed to HIV-ALF or healthy-ALF with an MOI of 10:1 for 2 h. Coverslips were fixed with 4% paraformaldehyde and permeabilized with cold methanol as described (27, 31). To assess *M.tb* intracellular trafficking, macrophage lysosomal compartments were stained for LAMP1 marker using a mouse anti-human LAMP1 antibody (H4AB; Developmental Hybridoma Bank, Iowa City, IA) overnight at 4°C, followed by staining with an APC-fluorescent anti-mouse secondary antibody (Molecular Probes, Eugene, OR) or with matched isotype controls for 60 min at 37°C. Coverslips were mounted as described above for confocal microscopy. Phagosome-containing *M.tb* was seen in green (GFP), late endosomal-lysosomal compartments in red (LAMP-1), and co-localization indicative of phagosome-lysosome (P-L) fusion in yellow. Co-localization between phagosomes containing GFP-*M.tb* and lysosomes containing LAMP1 marker was quantified by counting >150 events per coverslip.

To determine *M.tb* replication within macrophages, the reporter *M.tb* strain (SSB-GFP, *smyc’*::mCherry *M.tb* Erdman) was grown on 7H11 agar containing 50 μg/mL hygromycin B (32, 36). Macrophage monolayers on glass coverslips were infected with this *M.tb* strain pre-exposed to control- or HIV-ALF at an MOI of 1:1 for 72 h. Fixed coverslips were mounted as above and imaged by confocal microscopy. At least 300 bacteria were quantified per condition. Quantification of the SSB-GFP reporter was accomplished by counting the number of bacteria with and without SSB-GFP replication foci. All microscopy data were analyzed by using the Zeiss ZEN software.

For intracellular survival, MDM monolayers were infected with ALF-exposed *M.tb* (GFP, Red-Cherry tomato or Lux) with an MOI of 1:1 and cells were incubated for 2 h at 37°C, 5%CO_2_, as described (31, 33). Subsequently, infected MDM monolayers were washed with warm RPMI (at 37°C) and further incubated at 37°C, 5%CO_2_ in RPMI with 2% autologous human serum for the indicated times post-infection (2 h for association and 24 h to 144 h for survival and bacterial replication). *M.tb* intracellular survival was assessed by two approaches. For the luciferase assay, MDMs were infected with *M*.*tb*-Lux, and bacterial bioluminescence was measured in relative luminescence units (RLUs) every 24 h for up to seven days using a GloMax Multi Detection System (Promega) (35). For colony-forming unit (CFU) assays, infected MDMs at the indicated time period post-infection were lysed, diluted, and plated on 7H11 agar plates. The numbers of CFU were enumerated after bacterial growth for 3 to 4 weeks at 37°C (27, 31, 41).

### *M.tb* growth and intracellular trafficking after human SP-D replenishment in ALF

*M.tb-*Lux was exposed to control- or HIV-ALF with or without (as control) supplemented human recombinant SP-D (5 μg/mL, R&D Systems). After exposure, ALF-exposed *M.tb* was washed to remove all traces of ALF and re-suspended in single bacterial suspensions using sterile 0.9% NaCl. MDMs were infected with ALF-exposed *M.tb* at an MOI of 1:1 for RLU growth assay and at an MOI of 10:1 for intracellular trafficking studies as described above.

### Statistical analysis

Statistical significance was determined using GraphPad 9.3.1 Prism software. In these studies, “n” values represent the number of times that an experiment was performed using different human donors for cells and different human donors for ALFs (for both control- and HIV-ALFs). Statistical analyses were performed using an unpaired, 2-tailed Student’s *t-*test to compare the difference between two groups, and One-way ANOVA to compare differences among multiple groups, followed by Tukey’s post-hoc analysis.

## RESULTS

### Levels of oxidized proteins are high in the alveolar environment of PLWH compared to control donors

First, we normalized the ALF samples based on their physiological phospholipid content within the lung as we described previously (21, 26–29, 31–33, 42, 43). To analyze the effects of oxidative stress in the lung environment of PLWH, we measured carbonyl and 3-nitrotyrosine protein modification in control- and HIV-ALFs, indicators of irreversible protein oxidation by ROS and RNS, respectively (**Fig 1A, B**) (21, 32, 33, 44). We also measured myeloperoxidase (MPO) levels, indicative of an oxidative environment partially driven by a neutrophil influx (**Fig 1C**). Our results indicate that HIV-ALF soluble proteins contained significantly higher levels of carbonyl and nitrotyrosine modifications relative to control-ALF. MPO levels trended downward in HIV-ALF but did not reach statistical significance.

**Figure 1.**
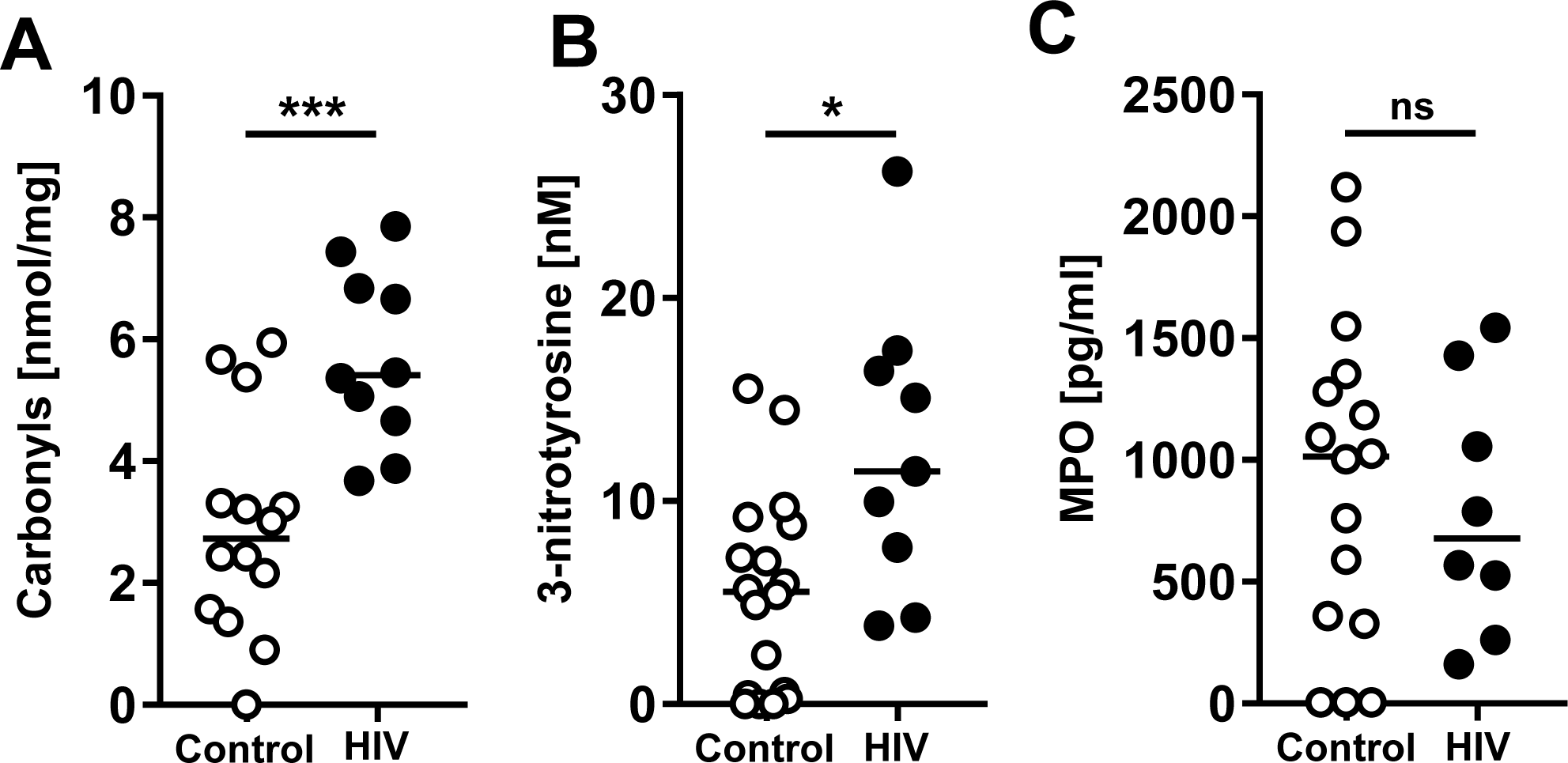
Oxidized protein levels are high in the alveolar environment of PLWH compared to control donors. Each dot represents ALF from an individual subject. ALFs from control (n= 14-16) and PLWH (HIV+, n= 8-10) subjects. (**A**) Protein carbonyl, (**B**) 3-nitrotyrosine residues were detected by ELISA, and (**C**) Myeloperoxidase (MPO) was detected by human multiplex Luminex assay. Unpaired Student’s *t* test, \**p*< 0.05; \*\*\**p*< 0.0005. Each sample corresponds to ALF obtained from different human donors.

### Immune mediators are altered in ALF samples from PLWH

Several ALF innate immune soluble components function as opsonic factors for microbes by directing them to specific phagocytic pathways in phagocytes. We opted to focus on the levels and functions of innate soluble proteins including components of the complement cascade (C2, C3, C5, and C9) and collectins, such as SP-A, SP–D, and mannose binding lectin (MBL), given their prominent innate role in the lung alveolar compartment and *M.tb* infection (45–47). We found that HIV-ALF samples had significantly lower levels of complement components, specifically C2, C3, and those involved in the formation of the membrane attack complex, C5 and C9 (**Fig 2A**). We also observed significantly lower SP-D levels in HIV-ALF, whereas SP-A and MBL levels in HIV-ALF were not significantly different from control-ALF (**Fig 2B**).

**Figure 2.**
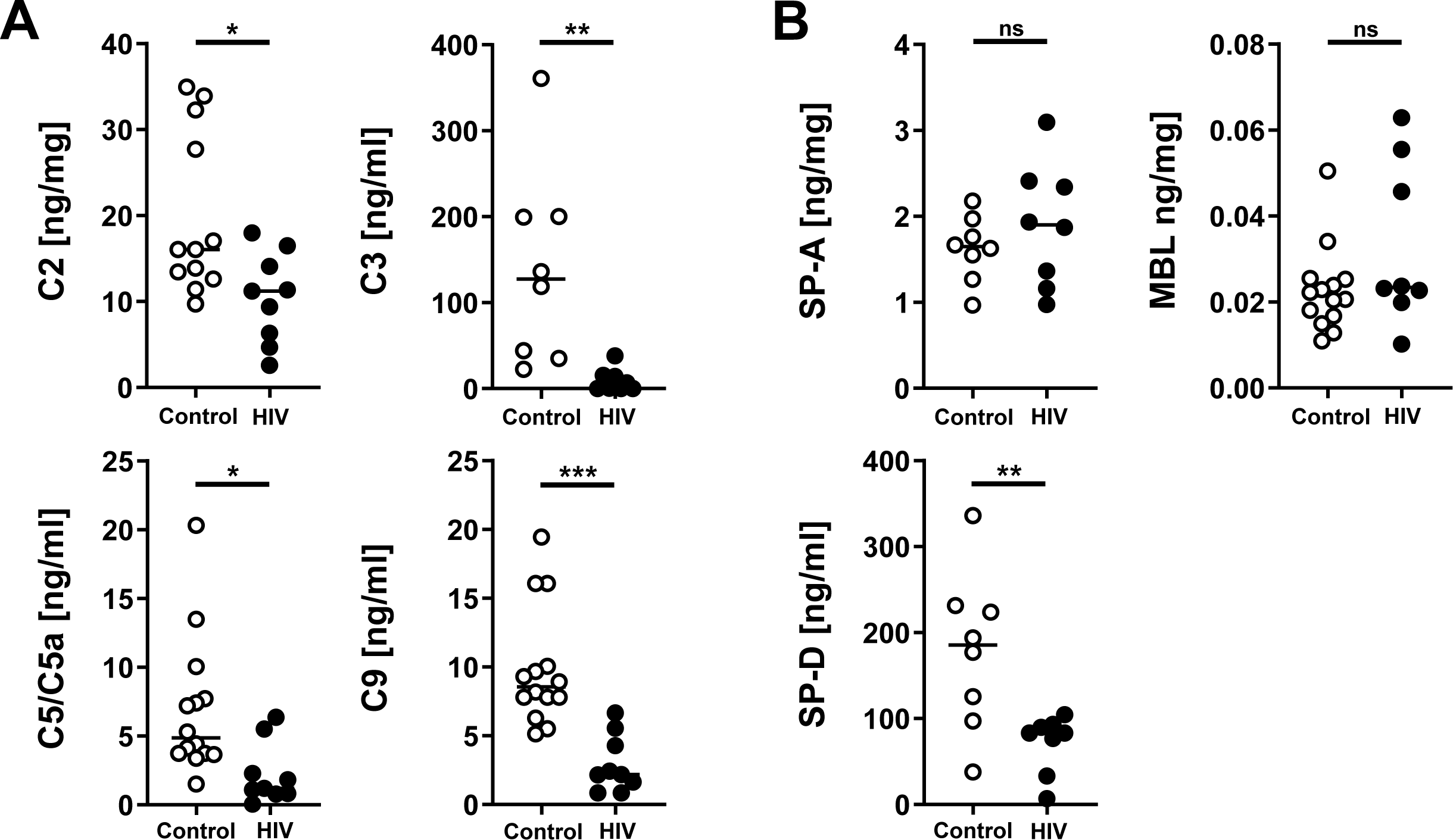
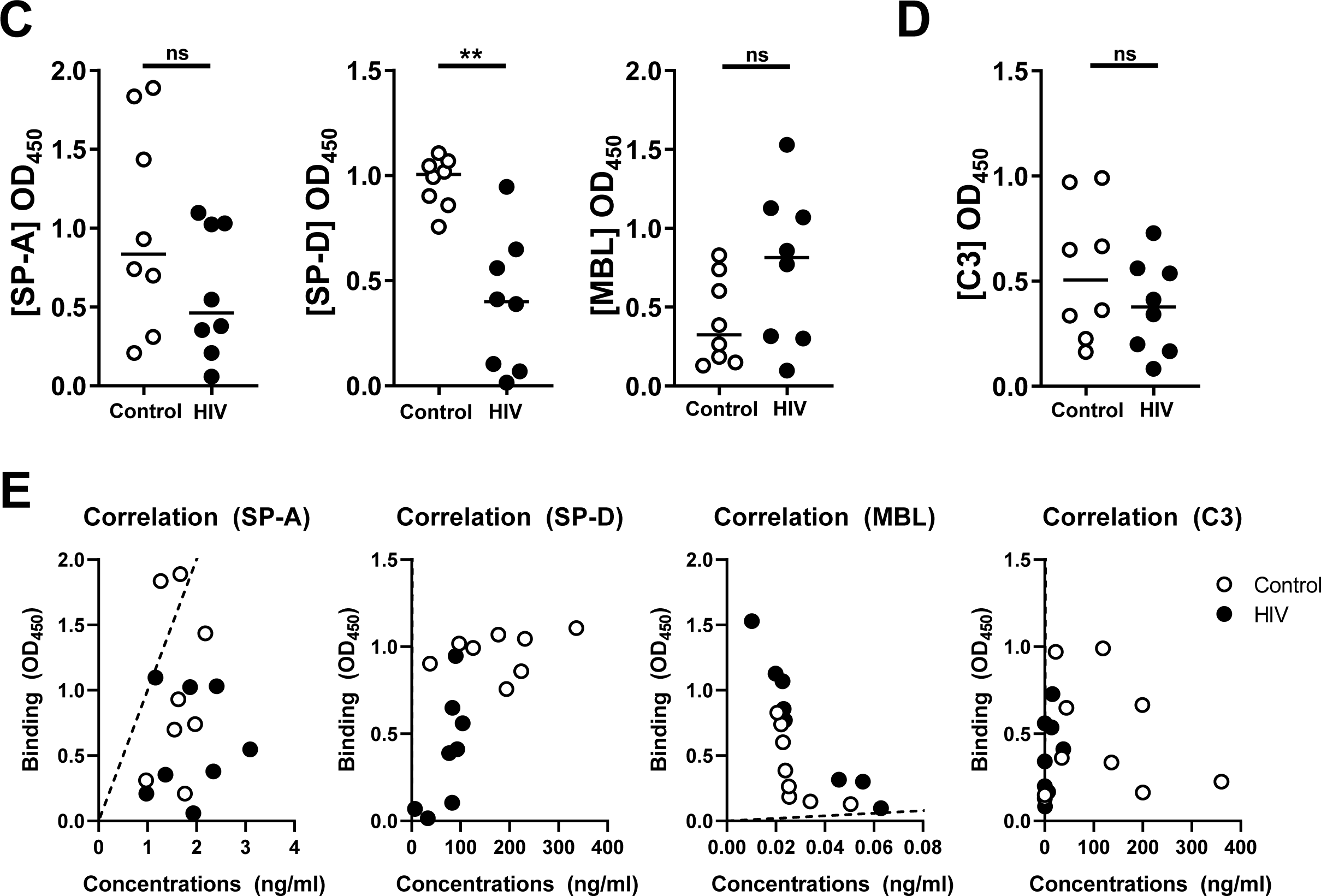
Measurement of innate soluble mediators in ALF of both PLWH and control individuals. Each dot represents ALF from an individual subject. ALFs from control donors (n= 8-17) and PLWH subjects (HIV+, n= 7-9). (**A**) Complement components: C2, C5/C5a, C9; and (**B**) collectins SP-D, MBL were detected by Luminex. Complement component C3 and collectin SP-A were detected by ELISA. *M.tb* Erdman strain single cell suspensions were exposed to control-ALF or HIV-ALF. Exposed-*M.tb* bacteria were washed, suspended in isotonic buffer, and plated onto 96-well plates. Monoclonal antibodies directed against (**C**) collectins SP-D, SP-A, MBL and (**D**) complement component C3 were used to determine their amounts bound to *M.tb*. For assay controls, purified SP-D, SP-A, MBL and C3 were used. Relative quantities of bound protein were quantified by standard ELISA by measuring the absorbance at OD_450_. (**E**) Correlations between the concentration levels and binding of SP-A, SP-D, MBL, and C3. Notice that regression line for SP-D and C3 overlaps with the *y*-axis. Unpaired Student’s *t*-test, *p< 0.05; **p<0.005, ***p< 0.0005. Each sample corresponds to ALF obtained from different human donors.

We used an *M.tb*-binding assay to assess the functions of several of these innate immune proteins present in ALF, which play important roles as opsonins during *M.tb* infection (46, 47). SP-D binding to *M.tb* was significantly decreased in HIV-ALF, while SP-A, MBL, and C3 binding remained unaltered in HIV-ALF compared to control-ALF (**Fig 2C, D**). It is worth noting that although C3 levels were lower in HIV-ALF, C3 binding to *M.tb* was similar in HIV-ALF and control-ALF (**Fig 2A, D**). In contrast, SP-D had both lower levels and lower binding in HIV-ALF when compared to control-ALF (**Fig 2B, C**). In both control-ALF and HIV-ALF, the amount of SP-A, SP-D, MBL and C3 that bound to *M.tb.* did not directly correlate with their concentrations (**Fig. 2E**), indicating that the observed lower binding of SP-D is not solely due to its lower level.

### The ALF proteome is altered in PLWH

We performed a proteomic assessment of both HIV-ALF and control-ALF by data-independent acquisition mass spectrometry. To identify differentially abundant proteins (DAPs) of interest, we calculated their log2 fold-changes in the HIV-ALF relative to those in control-ALF. In general, HIV-ALF contained higher levels of inflammation-associated proteins (**Fig 3A, Table S1**), and lower levels of anti-oxidation, complement component, and surfactant proteins (**Fig 3B, D and Table S1, S2**). However, the relative levels of antimicrobial proteins (**Fig 3C, Table S2**), immunoglobulins (IG) and IG receptor proteins (**Fig 3E, Table S3**), and hydrolytic proteins (**Fig 3F, Table S3**) in HIV-ALF did not follow this pattern; indeed, they showed distinct differential abundance. These results indicate that HIV infection seems to alter global protein abundance in the alveolar airspace. Of note, a major antibody of the lung mucosa, IgA (composed of IgA heavy chain alpha IgHA1 and its analog IgHA2) was detected at a lower abundance in the HIV-ALF (**Fig 3E**). IgJ, which is required for IgA and IgM secretion into the lung mucosa (48), was also detected at lower levels in HIV-ALF (**Fig 3E**).

**Figure 3.**
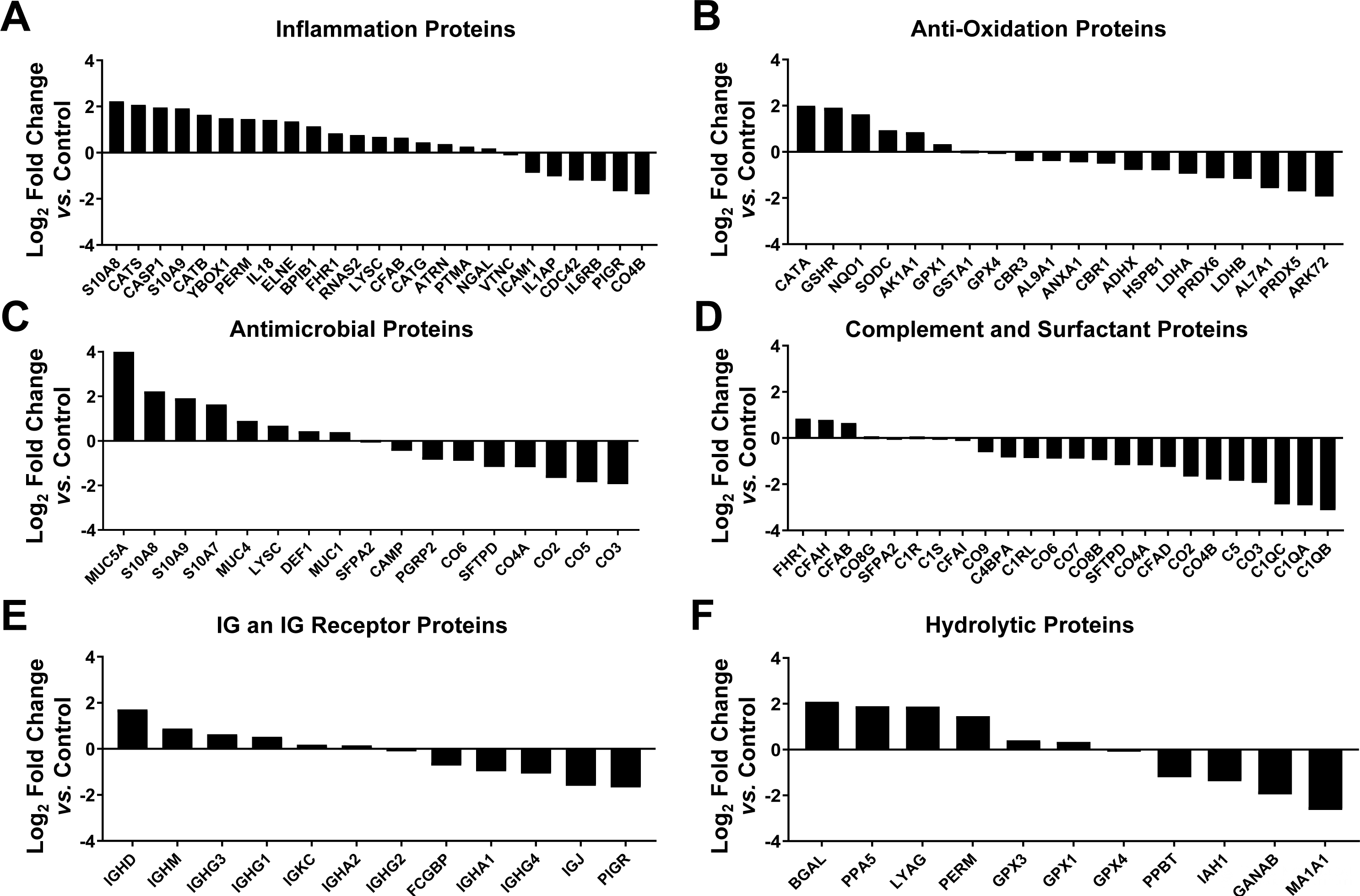
Proteomic analyses of ALF of both PLWH and control individuals. Six major categories of ALF-derived proteins were identified: (**A**) inflammation proteins, (**B**) antioxidation proteins, (**C**) antimicrobial proteins, (**D**) complement and surfactant proteins, (**E**) IgG & IgG receptor proteins, and (**F**) hydrolytic enzymes or hydrolases. Differentially abundant proteins (DAPs) were identified by calculating their log2 fold-changes in the HIV-ALF relative to those in control-ALF. Each sample corresponds to ALF obtained from different human donors. For names of the proteins depicted and statistics, see supplemental material and Tables S1 to S3.

### Exposure of *M.tb* to HIV-ALF provides a bacterial association and growth advantage in primary human macrophages

To test our hypothesis that HIV-induced oxidative stress in the lung alveolar environment results in dysfunctional soluble innate responses predisposing PLWH to a higher risk of respiratory infections including *M.tb*, we determined if exposure of *M.tb* to HIV-ALF influences the capacity of human macrophages (MDMs) to control *M.tb* infection. First, we assessed whether HIV-ALF had an impact on the capacity of macrophages to recognize *M.tb*. Control- and HIV-ALF-exposed *M.tb* were used to infect MDM monolayers and bacterial-cell association determined. Our results indicate that HIV-ALF-exposed *M.tb* has a significantly higher association with MDMs when compared to control-ALF-exposed *M.tb* (**Fig 4A**, supplemental **Fig S1A**).

**Figure 4.**
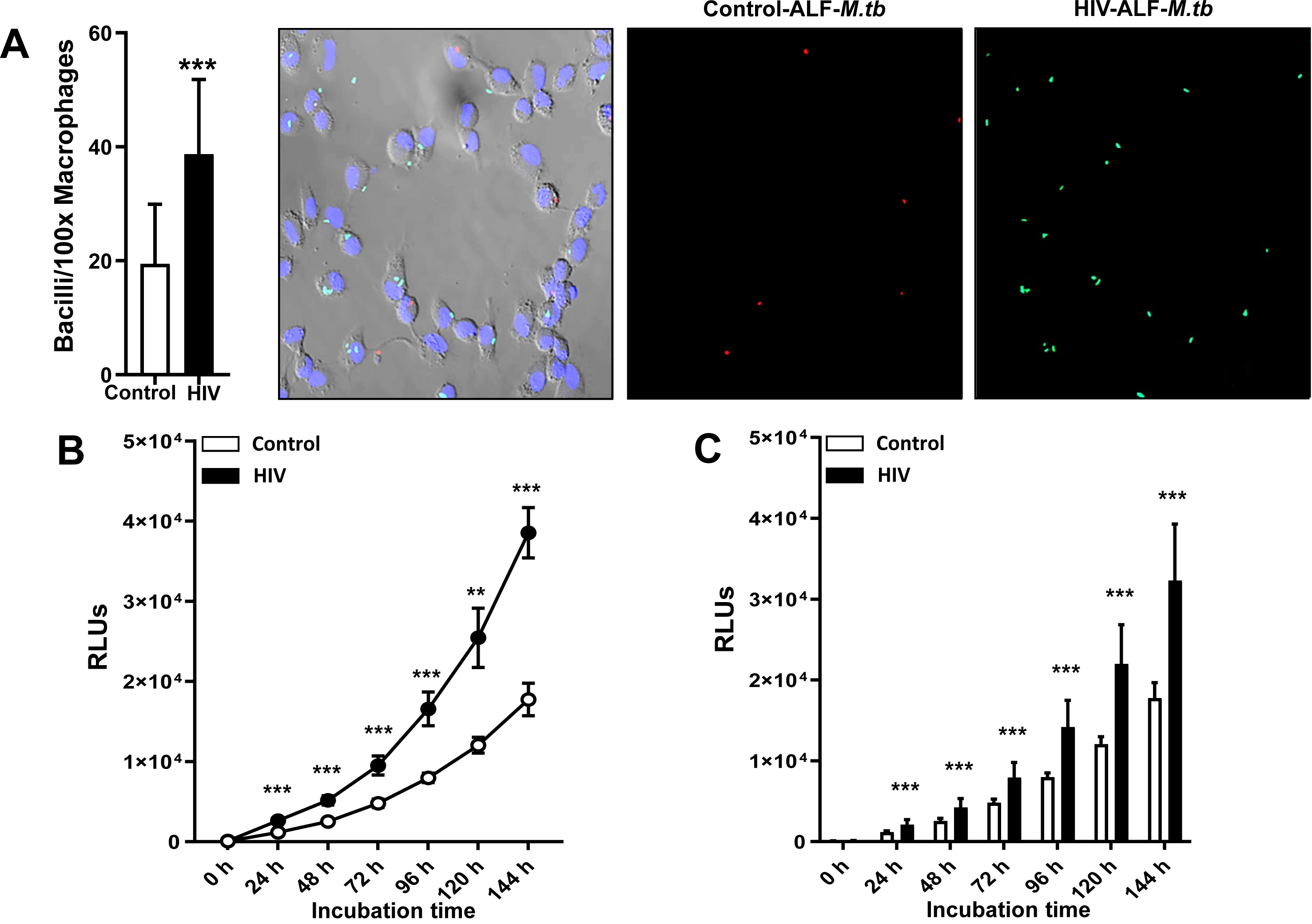
PLWH ALF favors bacteria-cell association and increased *M.tb* growth within human macrophages *in vitro*. (**A**) GFP-*M.tb* Erdman and Red-Cherry tomato *M.tb* Erdman bacteria were exposed to HIV-ALF or control-ALF, respectively. After exposure, these bacteria were mixed at 1:1 ratio, and macrophages on coverslips were infected with an MOI 10:1 of the bacterial mixture (n= 3/group). Coverslips were processed for and analyzed by confocal microscopy. Shown are GFP-*M.tb* Erdman exposed to HIV-ALF (Green) and Red-Cherry tomato *M.tb* Erdman exposed to control-ALF (red). Graph shows percentage of bacterial association with macrophages infected with *M.tb* exposed to healthy-ALF or HIV-ALF. Counted >150 macrophages per coverslips. Unpaired Student’s *t*-test; Healthy *vs* HIV+; \*\*\**p*<0.0005. (**B**) *M.tb* H_37_R_v_-Lux was exposed to HIV-ALF or control-ALF. Macrophage monolayers were infected with ALF-exposed *Mtb*-Lux (MOI 1:1) and *M.tb* intracellular growth was assessed by measuring bacterial bioluminescence in terms of RLUs at different time points, representative experiment and (**C**) Overall data from n= 4. ANOVA Tukey-Posttest; Healthy *vs.* HIV+; **p<0.005; ***p<0.0005. Each sample corresponds to ALF obtained from different human donors.

For evaluate intracellular growth of HIV-ALF-exposed *M.tb vs.* control-ALF-exposed *M.tb*, we employed both bacterial RLU (**Fig. 4B, C**) and traditional CFU (supplemental **Fig S1B**) assays in MDMs by using a luciferase-expressing strain *M.tb*-Lux and GFP-expressing *M.tb* Erdman, respectively (27, 31, 33, 35). Although we observed a significant difference in association (**Fig. 4A**), the uptake (quantity of intracellular bacilli at 2 h) was similar (**Fig 4B, C** and supplemental **Fig S1B**). Despite this similar uptake, *M.tb* exposed to HIV-ALF showed significantly increased growth in MDMs compared to *M.tb* exposed to control-ALF, which became more apparent over time (although we note that the luciferase and CFU assays are less sensitive than the association assay by microscopy) (**Fig 4B, C**).

### Exposure of *M.tb* to HIV-ALF increases bacterial replication rate and reduces its P-L fusion in primary human macrophages

To investigate the characteristics of the growth of HIV-ALF-exposed *M.tb* in MDMs, we evaluated *M.tb* intracellular replication and intracellular trafficking. Using a fluorescent *M.tb* replication reporter strain (SSB-GFP, *smyc’*::mCherry *M.tb*) (32, 33, 36), the number of *M.tb* bacilli with SSB-GFP foci were counted and then the percentages calculated to determine the rate of active replication. Our results indicate that HIV-ALF-exposed *M.tb* had a small but significantly increased replication rate compared to control-ALF-exposed *M.tb* (control: 13.62 ± 1.48% *vs.* HIV: 22.28 ± 0.68%) (**Fig 5A, B**).

**Figure 5.**
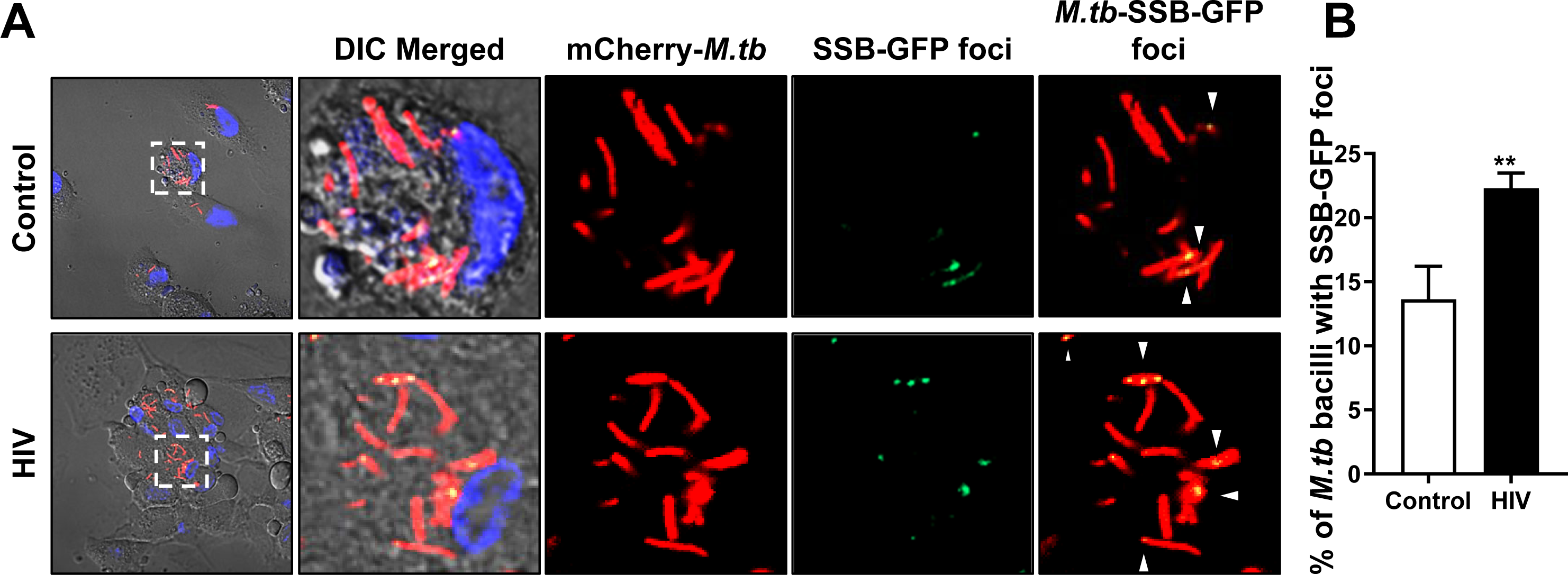
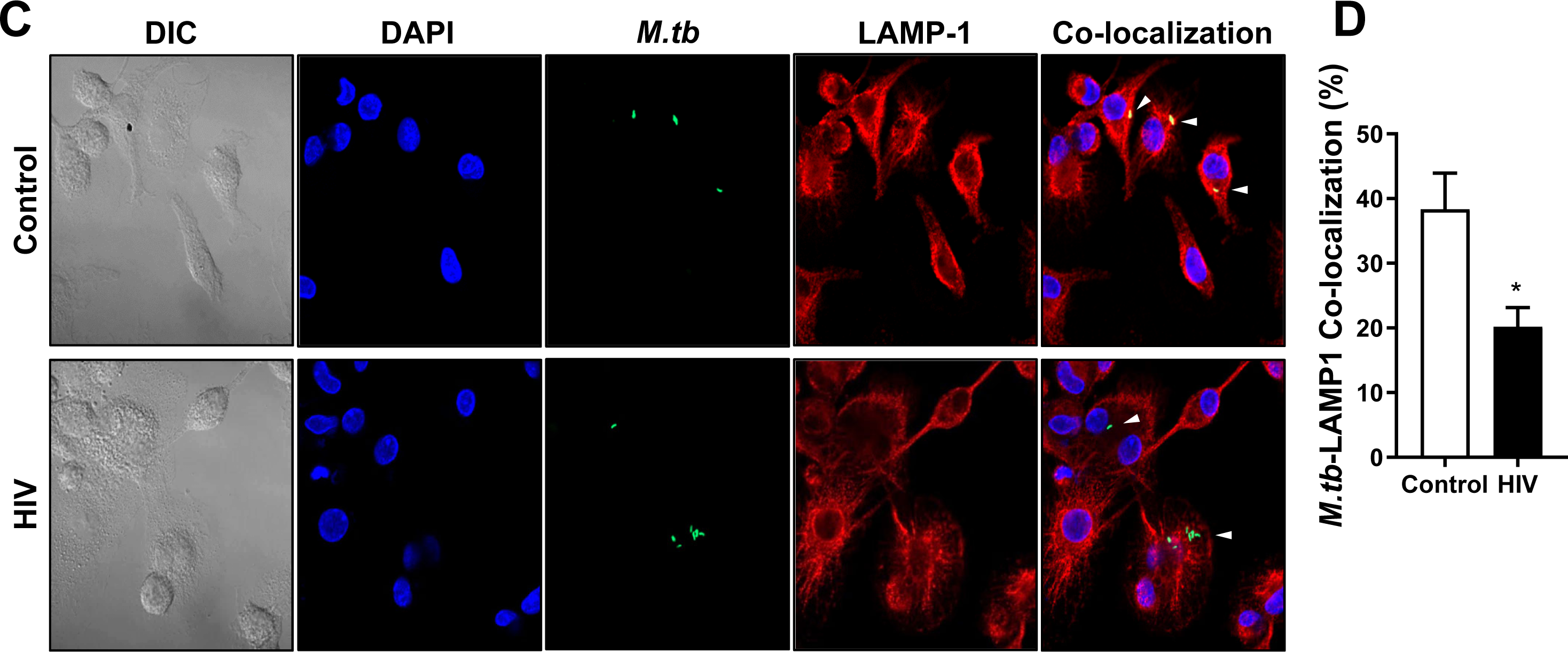
Assessment of intracellular bacterial replication and P-L fusion events in human macrophages containing *M.tb* pre-exposed to healthy-ALF or HIV-ALF. (**A**) Reporter fluorescent strain SSB-GFP, *smyc’*::mCherry *M.tb* was exposed to HIV-ALF or control-ALF prior to infection of human macrophages on coverslips for 72 h and *M.tb* replication rate was determined by confocal microscopy. Shown are representative confocal images of *M.tb*-infected macrophages on left panels (top and bottom), where region indicated by white dashed-line is shown in large magnification on right with control-ALF exposed samples on top panels and HIV-ALF exposed samples on bottom panels. SSB-GFP+ve *M.tb* are indicated by white arrowheads, showing merged (yellow) foci. (**B**) Percentage of SSB-GFP-positive *M.tb* exposed to control-ALF or HIV-ALF quantified by counting >150 events per coverslip (n= 3). Unpaired Student’s *t*-test; HlV-ALF *vs.* Control-ALF; **p<0.005. (**C**) GFP-*M.tb* Erdman was exposed to HIV-ALF or control-ALF prior to infection of human macrophages on coverslips for 2 h (n=3). P-L fusion events were visualized with confocal microscopy and enumerated by counting at least >150 independent events per coverslip. Shown are DIC, macrophage nucleus (DAPI, blue), phagosome containing GFP-*M.tb* (green), lysosomes (LAMP-1+, red), and P-L fusion events (co-localization, yellow). (**D**) Graph shows the percentage of co-localization of LAMP-1 (lysosomal marker) with phagosomes containing *M.tb*. Unpaired Student’s *t*-test; Healthy *vs* HIV; n=3 *p<0.05; **p<0.005. Each sample corresponds to ALF obtained from different human donors.

To measure intracellular trafficking, we quantified the number of *M.tb* phagosomes fused with lysosomes (LAMP-1+) and expressed the result as a percentage of *M.tb*-LAMP-1 co-localization. It is known that *M.tb* phagosomes have limited fusion with lysosomes.(49) Our results show that there was less co-localization with lysosomes by phagosomes from HIV-ALF-exposed *M.tb* than control-ALF-exposed *M.tb* (HIV: 20.2 ± 1.7% *vs.* control: 38.37 ± 37%) (**Fig 5C, D**). Collectively, our data indicate that a higher replication rate and significantly reduced P-L fusion contribute to the increased growth of HIV-ALF-exposed *M.tb* in MDMs.

### Supplementation of HIV-ALF with SP-D restores the macrophage capacity to control HIV-ALF-exposed *M.tb* infection

SP-D binds avidly to the surface of *M.tb* via lipoarabinomannan (23), driving an increase in P-L fusion of SP-D-coated *M.tb*, resulting in a better control of *M.tb* growth in MDMs (24). We supplemented HIV-ALF with SP-D to test the hypothesis that the increased *M.tb* growth observed in MDMs infected with HIV-ALF-exposed *M.tb* (**Fig 4B, C**) is due mainly to the low levels and low binding of SP-D in HIV-ALF compared to control-ALF (**Fig 2B, C**). Subsequently, *M.tb* bacilli (*M.tb*-Lux for growth assay and GFP-*M.tb* for P-L fusion assay) were exposed to SP-D**-**supplemented HIV-ALF and control-ALF prior to infection of human MDMs (33). Results showed that the addition of SP-D to the HIV-ALF allowed macrophages to regain control of the infection at the same levels that were observed for control-ALF-exposed *M.tb* (**Fig 6A, B**), and that this effect is via increasing P-L fusion events to reach the same levels as those observed in MDMs infected with control-ALF–exposed *M.tb* (a 17.6% increase in P-L fusion) (**Fig 6C, D**). These data support the concept that the binding of SP-D to *M.tb* is impaired in HIV-ALF and that its supplementation restores the capacity of human macrophages to control HIV-ALF-exposed *M.tb* growth.

**Figure 6.**
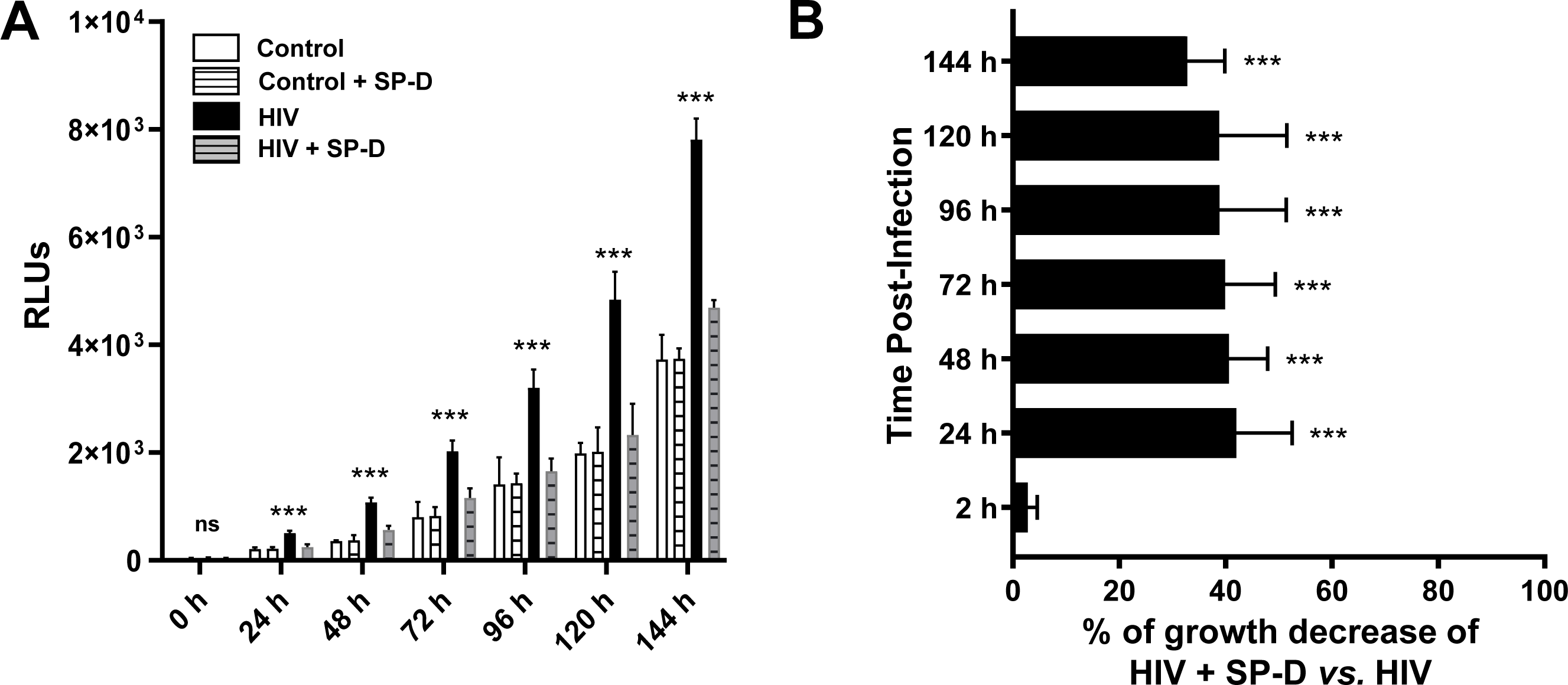
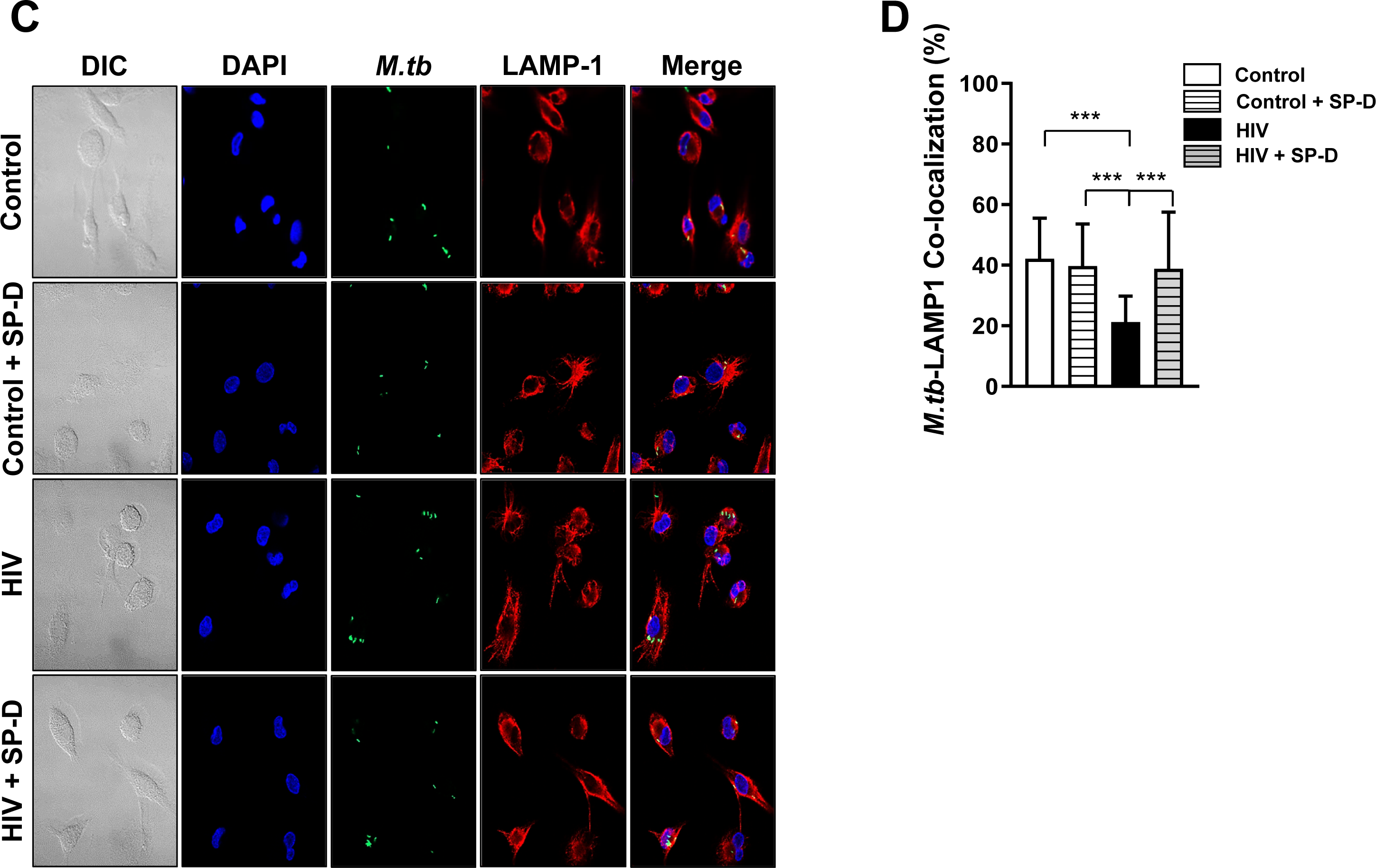
Addition of SP-D to HIV-ALF restores macrophage control of *M.tb* growth. (**A**) *M.tb* H_37_R_v_-Lux was exposed to HIV-ALF, control-ALF, or to HIV-ALF or control-ALF each being supplemented with SP-D at its physiological concentration. Human macrophages were infected with these differently-exposed *M.tb* and bacterial intracellular growth was measured in RLUs at different time points through 144 h. Representative experiment of n= 3, using ALF and macrophages in each case from different human donors. ANOVA Tukey-Posttest; HIV *vs.* Control; ***p<0.0005. (**B**) Overall data (n=3) showing % of growth decrease for HIV-ALF + SP-D exposed *M.tb vs.* HIV-ALF exposed *M.tb*. ***p<0.0005. (**C**) GFP-*M.tb* Erdman was exposed to HIV-ALF, control-ALF, HIV-ALF+SP-D or control-ALF+SP-D as described above prior to infection of macrophages on coverslips for 2 h (n= 3). P-L fusion events were visualized with confocal microscopy and enumerated by counting at least >150 independent events per coverslip. Shown are DIC, macrophage nucleus (DAPI, blue), phagosome containing GFP-*M.tb* (green), lysosomes (LAMP-1+, red), P-L fusion events (co-localization, yellow). (**D**) Graph shows percentage of P-L fusion in terms of *M.tb*-LAMP-1 co-localization. Unpaired Student’s *t*-test; Healthy *vs* HIV+; n= 3 ***p<0.0005. Each sample corresponds to ALF obtained from different human donors.

### Immune responses are altered in the alveolar space of PLWH

Peripheral inflammation is known to persistently occur in PLWH (10, 11). Thus, we examined whether this also occurs in the lung environment, which is normally tightly immune-regulated (50). Our results indicate that Th1 cytokines IFNγ, TNF, IL-12 and IL-2; Th2 cytokines IL-4, IL-10 and IL-13; and Th17 cytokines IL-17 and IL-21 were significantly lower in HIV-ALF compared to control-ALF, but IL-6, IL-22, and IL-27 were unchanged between control- and HIV-ALF (**Fig 7A-C**). HIV-ALF also had significantly lower levels of the growth factor GM-CSF and the inflammation marker TREM-1 (**Fig 7D**). Conversely, chemoattractants such as IL-8 and CCL2 were significantly increased in HIV-ALF (**Fig 7E**). Taken together, HIV-ALF had lower levels of inflammatory mediators but increased levels of chemoattractants when compared to control-ALF.

**Figure 7.**
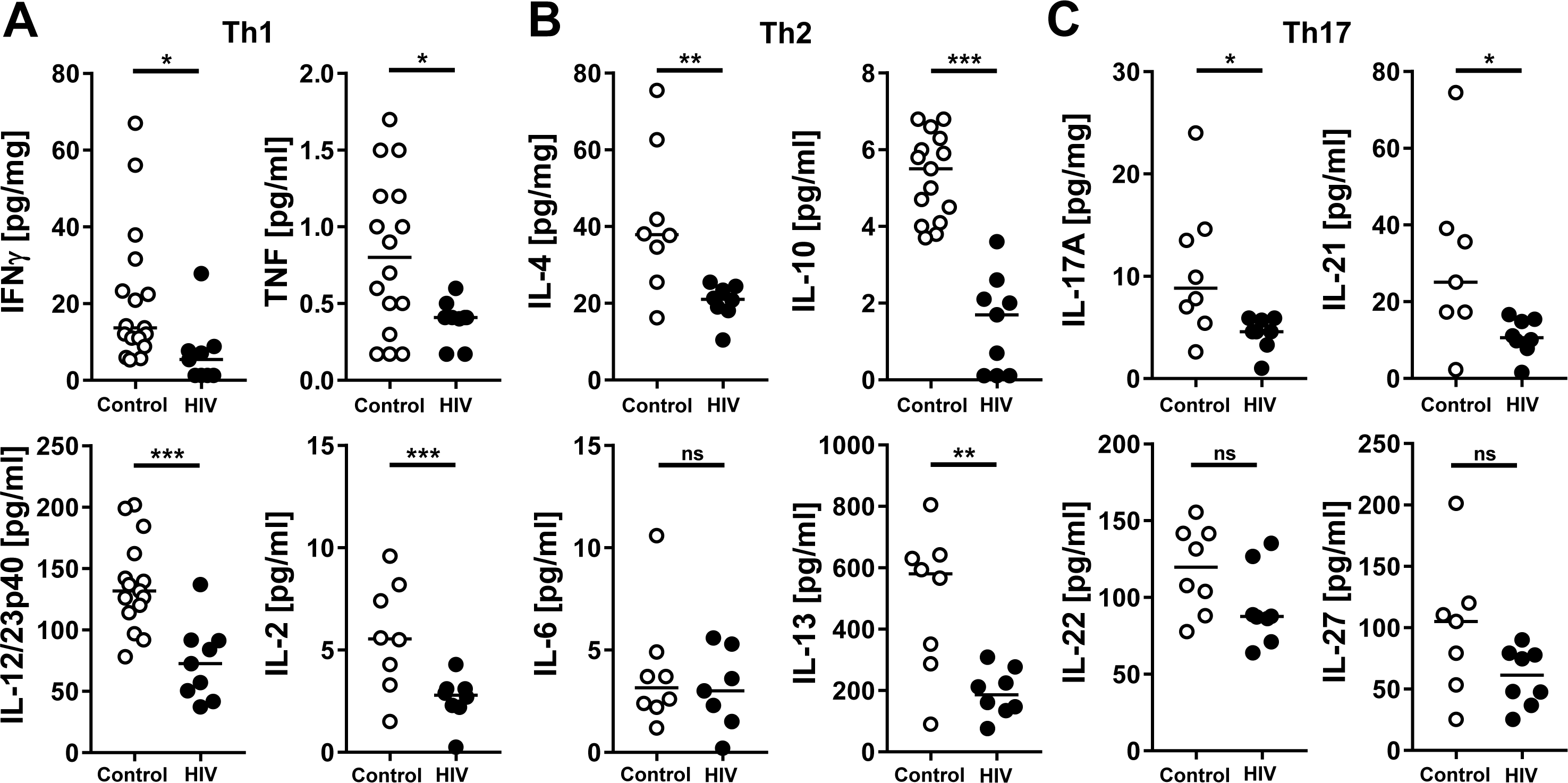
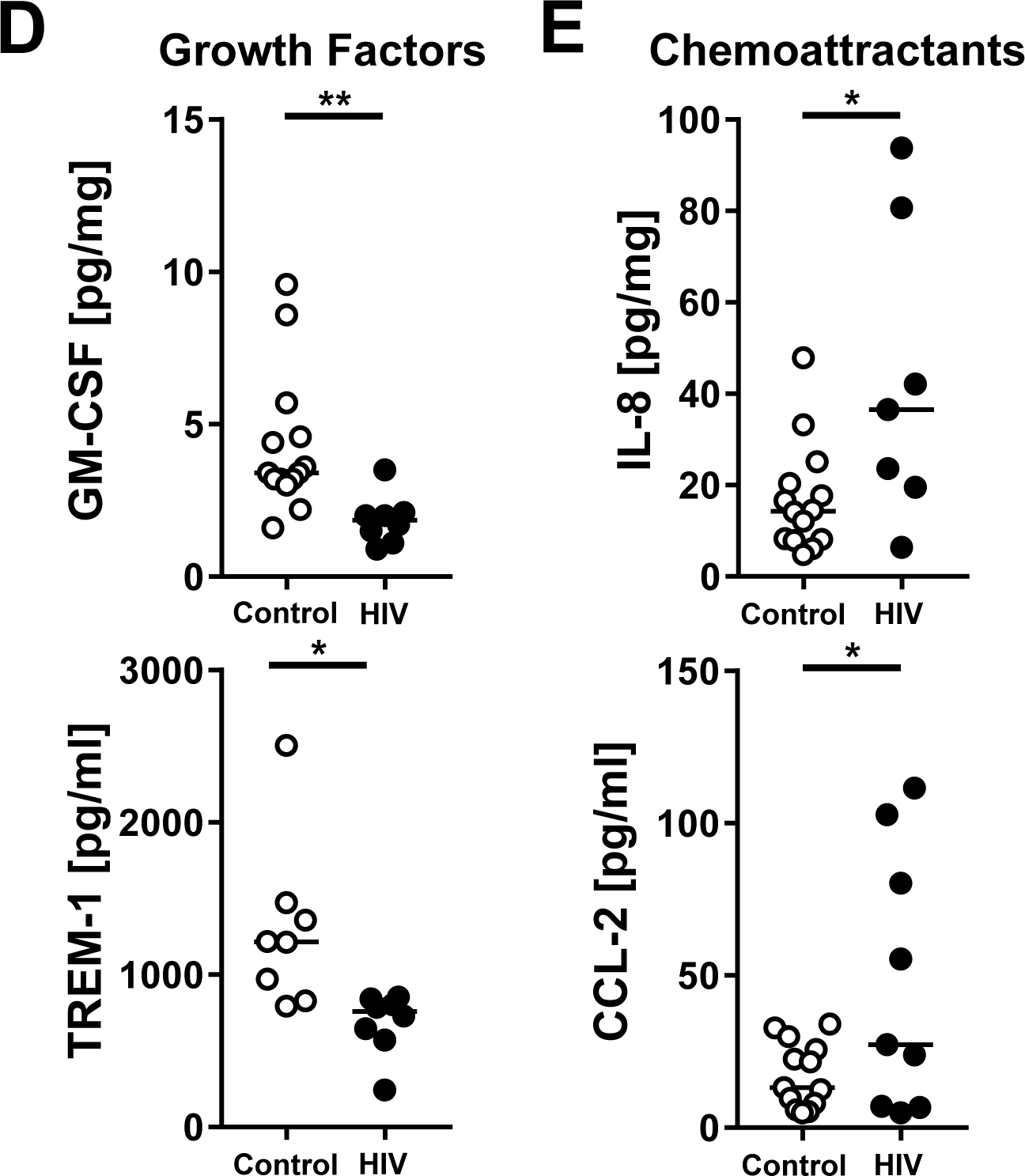
Measurement of cytokines, growth factors and chemokines present in ALF samples of both PLWH and control individuals. Each dot represents ALF from an individual subject. ALFs from healthy donors (n= 8-17) and HIV+ subjects (without ART; n= 7-9) were normalized by protein content (10 μg/well, by BCA). (**A**) Th1, (**B**) Th2, and (**C**) Th17 cytokines; (**D**) Growth factors TREM-1 and GM-CSF; and (**E**) Chemoattractants IL-8 and CCL-2 were measured by Human multiplex LUMINEX assay. Unpaired Student’s *t*-test, *p< 0.05; **p<0.005, ***p< 0.0005. Each sample corresponds to ALF obtained from different human donors.

## DISCUSSION

PLWH have a 25-fold increased risk of respiratory tract infections (51–53); even modest insufficiency of the immune system in the lung makes PLWH more vulnerable to microbial infections (51, 52), as well as, to other chronic diseases. In fact, *M.tb* is a frequent respiratory tract infection among PLWH and TB is the major cause of death in PLWH.

In this study, we demonstrated that PLWH have an alveolar environment marked by a high degree of protein oxidation and decreased levels of anti-oxidation proteins. This is accompanied by a decrease in Th1/Th2/Th17 responses, as well as lower levels of complement proteins and SP-D. In the case of SP-D, its lower levels did not correlate with its lower binding to *M.tb*. In this regard, SP-D binding to *M.tb* leads to bacterial aggregation and reduced bacterial growth in macrophages by increasing P-L fusion (24).

In this context, a role has been reported for some ALF components during HIV infection (51). SP-A and SP-D can modulate HIV infection by inhibiting infectivity of CD4 T cells or by stimulating HIV transfer from dendritic cells (DCs) to CD4 T cells (51). SP-D binds to HIV envelope glycoprotein 120 (gp120), blocking HIV binding and entry into macrophages (54). Our results indicate that HIV-ALF contains low levels of SP-D with decreased binding to microbes. These SP-D alterations may be a contributing factor for the increased susceptibility of PLWH to respiratory infections. Our *in vitro* studies with *M.tb* support this concept since supplementation of HIV-ALF with SP-D restored the ability of macrophage to control *M.tb* growth. These *in vitro* studies suggest that the low levels of SP-D and its decreased binding are major contributors to the altered innate immune control of *M.tb* during HIV infection although its role *in vivo* remains to be determined. Reduced levels and reduced binding of SP-D in the lungs may correlate with negative pulmonary outcomes in PLWH without ART (51). This is the case for other lung diseases such as pulmonary fibrosis, where lower levels of SP-D correlate with increased mortality (55). In contrast to SP-D, SP-A levels and binding to *M.tb* in the PLWH ALF did not significantly differ from those in control individuals. SP-A can regulate the macrophage response to infection in that SP-A facilitates the transfer of HIV from dendritic cells (DCs) to CD4 T cells (56, 57). Decreased binding of SP-D and unaltered SP-A may favor establishment of *M.tb* infection in lung macrophages. Conversely, SP-A promotes attachment of *M.tb* to macrophages by increasing surface exposure of the mannose receptor (CD206) on macrophages (58), which is increased during HIV infection driving IL-6 production. In this regard, while we observed an overall decrease in Th1/Th2/Th17 cytokine levels in the lungs of PLWH while IL-6 remained unaltered. This decrease in cytokine levels in HIV-ALF is also observed in *in vitro* experiments showing that HIV infection impairs the production of TNF by macrophages (59, 60). Thus, HIV infection may weaken the lung immune response against subsequent microbial infections because of dysregulation of inflammatory responses in the lung, generating an environment that favors pulmonary microbial infections (51). Our model is illustrated in **Fig 8**.

**Figure 8.**
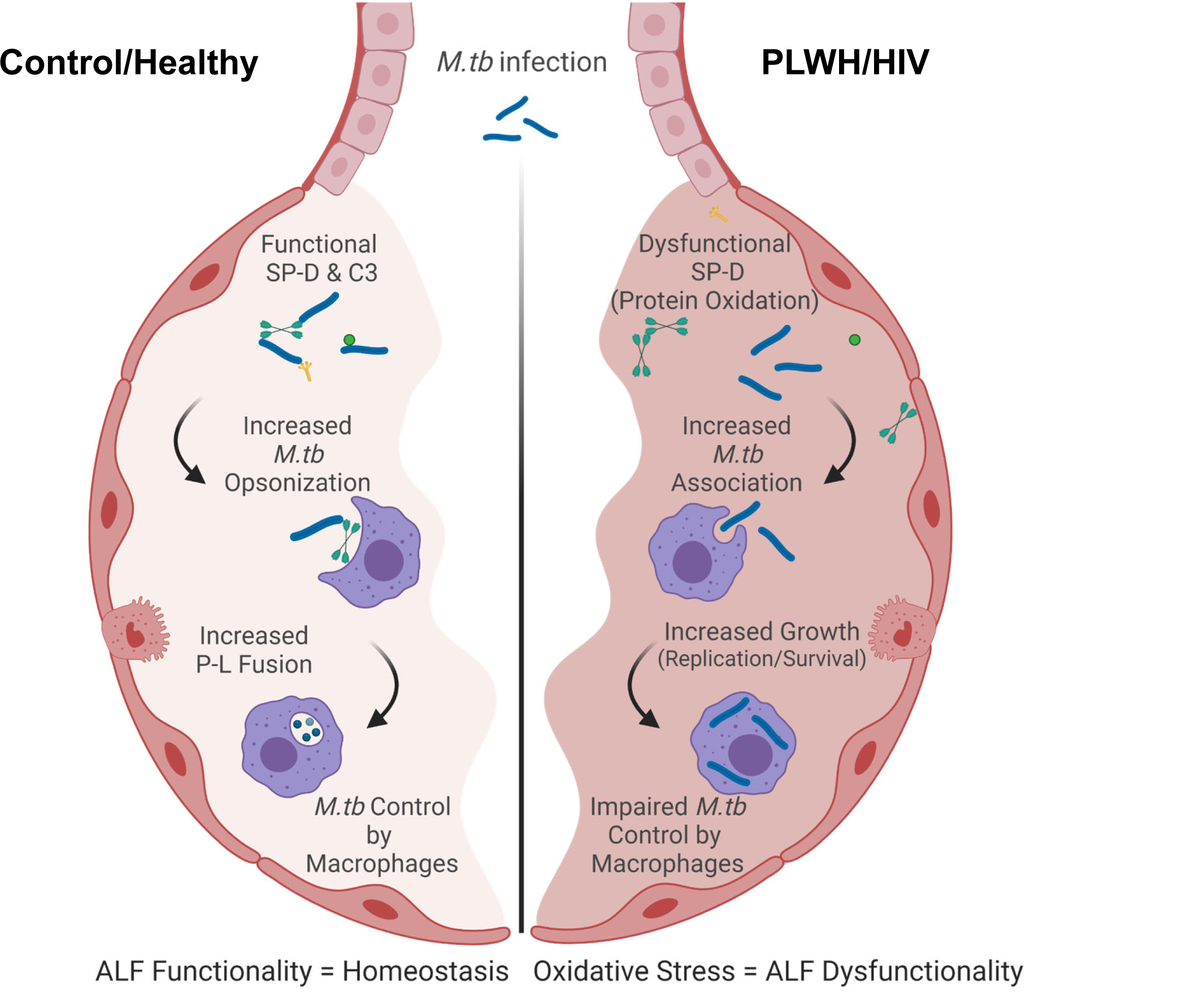
Model of the host responses to *M.tb* in the lung of PLWH compared to control. Local oxidative stress is detected in the lung of PLWH. This oxidative stress may drive the dysfunction of soluble immune components and inflammation in the lungs marked by decreased levels of Th1/Th2/Th17 cytokines. Lower levels of complement and SP-D are also detected in PLWH leading to their lower capacity to opsonize and/or modify the *M.tb* bacterial cell surface prior contacting host cells. This drives better recognition of HIV-ALF exposed *M.tb* by macrophages. Phagocytosed HIV-ALF exposed *M.tb* bacilli grow faster within macrophages, by further impairing P-L fusion and replicating faster within phagosomes. *In vitro*, this is related to SP-D function, since addition of excess of SP-D to HIV-ALF restores the capacity of macrophages to control HIV-ALF-exposed *M.tb*; thus SP-D plays an important role in maintaining lung homeostasis and innate immune activity against lung respiratory infections. Created in *BioRender.com*.

Previous studies have shown that alveolar macrophages in PLWH are functionally impaired (51, 61–64), with a reduced capacity for phagocytosis (65, 66). However, components in HIV-ALF may not be responsible for this observed effect, as exposure of *M.tb* to HIV-ALF resulted in an increase of *M.tb* association with macrophages obtained from control individuals.

Although the complement system is systemically activated by HIV infection (67), within the HIV-ALF of PLWH we observed low levels of the classical complement proteolytic cascade, including both early complement components (e.g., C1qa, C1qc) and terminal factors (e.g., C5b-9) involved in the formation of the membrane attack complex. C1q also binds to several receptors on myeloid cells, regulating their functions, including phagocytosis (68). C1q deficiency is associated with increased infections and autoimmunity, and C5 and C7 deficiencies are linked to *M.tb* pathogenesis (68–71). The role of complement in *M.tb* pathogenesis remains unclear, although C3 binding to *M.tb* cell envelope components leads to *M.tb* recognition by the complement receptor 3 (CR3) on human macrophages (22, 72). The outcome of this interaction in terms of *M.tb* control is unknown. Our results indicate that PLWH have significantly less C3 than HIV-negative individuals, which has implications for the establishment of *M.tb* infection in PLWH. Regarding the complement lectin pathway, MBL binds the gp120 on HIV and *M.tb* (67, 73–75); however, MBL levels and binding remained unaltered in the HIV-ALF of the PLWH studied.

We also found lower levels of IgJ and pIgR in HIV-ALF. IgJ is involved in the secretion of IgA and IgM into the lung mucosa. It is essential for the binding of polymeric immunoglobulin to pIgR, which forms a secretory component, facilitating the transcytosis of soluble polymeric isoforms of IgA and immune complexes from basal to the apical sides of alveolar epithelial cells (76). Deficiencies of IgJ and pIgR in HIV-ALF could contribute to the decrease in IgA levels observed in HIV-ALF, detected as lower levels of IgA heavy chain alpha, IgHA1 and its analog IgHA2. IgA deficiency in the lung could contribute to the susceptibility to respiratory infections observed in PLWH. Of relevance, IgJ plays a role in the activation of complement (77).

Our proteomics results confirm that HIV-ALF contains an array of inflammatory, antimicrobial, and oxidant-scavenging proteins that are upregulated [e.g., calprotectin (S10A8/S10A9) protein and mucins (MUC1, 4, and 5A), among others]. However, these could have altered functional consequences, as we observed for SP-D. Further studies are required to determine the impact of HIV infection on the overall physiological and immunological status of the lung, and how this influences the establishment of acute (e.g. SARS-CoV-2) and chronic (e.g. *M.tb*) respiratory infections. Our study does not address the impact of ART on the lung environment. Systemically, ART fails to restore full recovery of the innate and adaptive systems and does not reduce systemic inflammation (78). ART is known to improve the adaptive immune response in the lung; however inflammation persists despite the recovery of CD4 T cells and HIV viral suppression (51). ART also induces the immune reconstitution inflammatory system (IRIS) in the lungs, particularly influencing latent *M.tb* infection or failed TB treatment (51). Thus, the effects of ART on soluble innate immune components and their functions in the lung need to be evaluated further. We anticipate that HIV infection disturbs the oxidative status of the lung. This perturbation of the oxidative status of the lung may be partially restored by ART (79, 80), but irreversible levels of protein oxidation (e.g. irreversible modification of tyrosine to 3-nitrotyrosine by RNS) (44) will persist; thus, explaining why PLWH treated with ART still have a higher susceptibility to respiratory infections.

A potential limitation of this study is that PLWH participants declared that they were never or former smokers at the time of providing HIV-ALF samples. This was clinically recorded, trusting the participants’ responses. We normalized our ALF samples to the physiological phospholipid concentrations in the human lung. The actual status related to former smoking is not expected to influence our results, as published studies have shown that never- and former-smokers have similar ALF phospholipid content (34). It could be rationalized also that HIV infection drives lower levels of SP-D and thus its observed decrease in binding to *M.tb*. However, our data show that there is no direct correlation between SP-D levels and binding. Our results are in contrast with other studies conducted with a cohort in Malawi, where SP-D levels were higher in PLWH. However, the subjects in that study had AIDS with CD4 T cells <200, and SP-D binding was not assessed (81). A plausible explanation for our results is that the decrease in SP-D binding to *M.tb* could be explained by the presence of the soluble HIV Env gp120 subunit in HIV-ALF since HIV gp120 binds SP-D and could, thereby, block SP-D binding to *M.tb*. Alternatively, gp120 could also directly bind to *M.tb*. In these two scenarios, the presence of gp120 in HIV-ALF could redirect *M.tb* to a pathway favoring its intracellular replication and survival. However, neither explanation appears to be justified. First of all, we did not detect the presence of gp120 in HIV-ALF by MS analysis. And second, gp120 also binds to MBL (82), and no reduction in MBL binding to *M.tb* was observed in this study.

Overall, our data highlight the importance of lower levels and reduced binding of SP-D in PLWH that we posit plays a role in respiratory infection outcomes. The low inflammatory lung environment observed in PLWH differs from the high inflammatory lung environment that we observed in the elderly leading to their susceptibility to respiratory infections, including *M.tb* (21, 33, 83). In both cases, however, PLWH and the elderly, there is a high degree of oxidative stress in their lungs, leading to higher levels of oxidation of proteins in the innate immune system which significantly alters the host response and is associated with susceptibility to respiratory infections. Identifying the outcomes of HIV-associated lung oxidative stress on alveolar innate immune responses will be valuable for the development of host-directed therapies for PLWH and the elderly reducing their risk of respiratory infections.

## AUTHORS CONTRIBUTIONS

AA and JBT contributed to the design of the studies; AA, JIM, AKA, JMS, AOF, AGV, and STW, contributed to the experimental procedures, and gathering data, and data analyses. PTD, MAG and MDW, provided the ALF samples. JJE, LSS and STW provided experimental guidance and feedback on data analysis. AOF did the BioRender figure. AA and JBT wrote the manuscript. All co-authors provided comments/edits in the final version of this manuscript.

## ACKNOWLEDGEMENTS

We would like to thank all the PLWH that participated providing samples to this study. We would like to thank Drs. Joanne Turner and Mark A. Endsley for their careful review of this manuscript and/or guidance in the discussion, and Dr. Anna Allué-Guardia for her assistance with the HIV-1 protein database. Mass spectrometry analyses were conducted at the University of Texas Health Science Center at San Antonio (UTHSCSA) Institutional Mass Spectrometry Laboratory, with expert technical assistance of Sammy Pardo and Dana Molleur.

---

**Table 1.** Clinical Parameters of PLWH in this study.

| BAL ID | Weight (Pounds) | BMI | Sex | Age (years) | Viral copies | CD4 T cell counts | ART |
| --- | --- | --- | --- | --- | --- | --- | --- |
| 154 | 209 | 28.34 | M | 54 | 48 | 227 | No |
| 155 | U* | U* | F | 32 | 48 | 661 | No |
| 161 | U* | U* | M | 45 | 48 | 605 | No |
| 179 | 189 | 25.63 | M | 44 | 883 | 532 | No |
| 182 | 135 | 21.14 | M | 35 | 65083 | 251 | No |
| 186 | 157 | 24.59 | M | 29 | 58836 | 286 | No |
| 190 | 150 | 25.74 | F | 20 | 48 | 710 | No |
| 204 | 174 | 27.25 | M | 23 | 150187 | 301 | No |
| 210 | 181 | 26.73 | M | 43 | 9413 | 429 | No |
| 219 | 191 | 23.25 | M | 30 | 210504 | 125 | No |

\*Unknown; M: Male, F: Female; Age (years), Viral copies (per μl), CD4 T cell counts (per μl), ART: antiretroviral treatment

**Table 2.** Cell types in BALs from PLWH in this study.

| BAL ID | Total Cell Count ($\times 10^6$) | Mϕ Count ($\times 10^6$) | % Mϕ | % L | % N | % E | % Ep |
| --- | --- | --- | --- | --- | --- | --- | --- |
| 154 | 16.81 | 13.95 | 83 | 9 | 4 | 1 | 3 |
| 155 | U* | U* | U* | U* | U* | U* | U* |
| 161 | U* | U* | U* | U* | U* | U* | U* |
| 179 | 33.65 | 31.3 | 93 | 7 | 0 | 0 | 0 |
| 182 | 24.89 | 22.9 | 92 | 5 | 3 | 0 | 0 |
| 186 | 31.18 | 29 | 93 | 4 | 0 | 0 | 3 |
| 190 | 3.13 | 2.6 | 83 | 8 | 8 | 0 | 1 |
| 204 | 16.20 | 8.1 | 50 | 50 | 0 | 0 | 0 |
| 210 | 23.49 | 19.5 | 83 | 17 | 0 | 0 | 0 |
| 219 | 27.87 | 22.3 | 80 | 20 | 0 | 0 | 0 |

\*Unknown; M $\phi$ : Macrophage; L: Lymphocytes; N: Neutrophils; E: Eosinophils; Ep: Epithelial cells.

