## Supplemental Material for "HIV Infection impairs the Host Response to *Mycobacterium tuberculosis* Infection by altering Surfactant Protein D function in the Human Lung Alveolar Mucosa"

**Figure 3: Names of the proteins depicted in Figure 3.**

**In panel A**: S10A8 (protein S100-A8), CATS (cathepsin S), S10A9 (protein S100-A9), CATG (cathepsin G), PERM (myeloperoxidase), BPIB1 (BPI fold-containing family B member 1), RNAS2 (non-secretory ribonuclease), YBOX1 (nuclease-sensitive element-binding protein 1), ELNE (neutrophil elastase), IL18, (isoform 2 of interleukin-18), FHR1 (complement factor H-related protein 1), S10AC (protein S100-A12), CFAB (complement factor B), LYSC (lysozyme C), MMP9 (matrix metalloproteinase-9), ATRN (attractin), NGAL (neutrophil gelatinase-associated lipocalin), PA21B (phospholipase A2), PTMA (prothymosin alpha), TXNL1 (thioredoxin), IL1AP (interleukin-1 receptor accessory protein), CD166 (CD166 antigen), VTNC (vitronectin), ICAM1 (intercellular adhesion molecule 1), IL6RB (interleukin-6 receptor subunit beta), CDC42 (cell division control protein 42), PIGR (polymeric immunoglobulin receptor), TRADD (TNF receptor binding type 1), CO4B (complement C4-B), and PAIRB (plasminogen).

**In panel B**: CATA (catalase), GSHR (glutathione reductas), NQO1 (NAD(P)H dehydrogenase), AK1A1 (alcohol dehydrogenase [NADP(+)]), GPX1 (glutathione peroxidase 1), SODC (superoxide dismutase [Cu-Zn]), GSTA1 (glutathione S-transferase A1), GPX4 (phospholipid hydroperoxide glutathione peroxidase), CBR3 (carbonyl reductase [NADPH] 3), AL9A1 (4-trimethylaminobutyraldehyde dehydrogenase), ANXA1 (annexin A1), CBR1 (carbonyl reductase [NADPH] 1), ADHX (alcohol dehydrogenase), HSPB1 (heat shock protein beta-1), LDHA (L-lactate dehydrogenase A), A1AT (alpha-1-antitrypsin), LDHB (L-lactate dehydrogenase B), PRDX6 (peroxiredoxin-6), ARK72 (aflatoxin B1 aldehyde reductase member 2), and AL7A1 (alpha-aminoadipic semialdehyde dehydrogenase).

**In panel C**: MUC5A (mucin-5AC), S10A7-A9 (protein S100-A8 to A9), MUC4 (mucin-4), MUC1 (mucin-1), LYSC (lysozyme C), DEF1 (neutrophil defensin 1), SFPA2 (pulmonary surfactant-associated protein A2), CAMP (cathelicidin antimicrobial peptide), CO6 (complement component C6), PGRP2 (N-acetylmuramoyl-L-alanine amidase), SFTPD (pulmonary surfactant-associated protein D ), and CO2 to 5 (complement C2 to C5).

**In panel D**: CFAH (complement factor H), FHR1 (complement factor H-related protein 1), CFAB (complement factor B), CO8G (complement component C8), SFPA2 (pulmonary surfactant-associated protein A2), C1R (complement C1r), C1S (complement C1s), CFAI (complement factor I), CO9 (complement component C9), C4BPA (C4b-binding protein alpha chain), CO7 (complement component C7), CO6 (complement component C6 ), C1RL (complement C1r subcomponent-like protein), CO8B (complement component C8 ), CFAD (complement factor D), SFTPD (pulmonary surfactant-associated protein D), CO2 to C5 (complement C2 to C5), and C1QA to C (complement C1q subcomponent subunit A to C).

**In panel E**: LYAG (alpha-glucosidase OS), GLU2B (glucosidase 2), AGAL (alpha-galactosidase A), GANAB (alpha-glucosidase AB), ARSA (arylsulfatase), PPBT (alkaline phosphatase), MA2B1 (alpha-mannosidase), and MA1A1 (mannosyl-oligosaccharide 1,2-alpha-mannosidase).

**In panel F**: LYAG (alpha-glucosidase OS), GLU2B (Glucosidase 2), AGAL (alpha-galactosidase A), GANAB (alpha-glucosidase AB), ARSA (arylsulfatase), PPBT (alkaline phosphatase), MA2B1 (alpha-mannosidase), and MA1A1 (mannosyl-oligosaccharide 1,2-alpha-mannosidase).

**Supplemental Table S1. Inflammation and anti-oxidation proteins.** Gene ID, name and t-test values.

**Supplemental Table S2. Antimicrobial, and complement and surfactant proteins.** Gene ID, name and t-test values.

**Supplemental Table S3. IG and IG receptor, and hydrolytic proteins.** Gene ID, name and t-test values.

**Supplemental Fig S1. HIV-ALF exposed *M.tb* in macrophages. (A)** Association of HIV-ALF vs. Control-ALF-exposed *M.tb* with macrophages showing a differential association favoring recognition of HIV-ALF-exposed *M.tb* over control-ALF-exposed *M.tb*. **(B)** PLWH ALF leads to increased *M.tb* growth within human macrophages *in vitro*. GFP-*M.tb* Erdman was exposed to HIV-ALF or healthy-ALF. MDMs were infected (MOI 1:1) with ALF-treated GFP-*Mtb* and cell monolayers were lysed at different time points, and then plated to assess bacterial burden by CFU counts (n= 3). Data shown are from a representative experiment. ANOVA Tukey-Posttest; Healthy *vs*. HIV+; *p<0.05; **p<0.005; ***p<0.0005. Each “n” value is an independent experiment using macrophages and ALF from different human donors.

### Table S1

#### Inflammation Proteins

| Gene ID | Name | t test |
|---|---|---|
| S10A8 | Protein S100-A8 | 0.29 |
| CATS | Cathepsin S | 0.0012 |
| CASP1 | Caspase-1 | < 0.0001 |
| S10A9 | Protein S100-A9 | 0.32 |
| CATB | Cathepsin B | 0.0076 |
| YBOX1 | Nuclease-sensitive element-binding protein 1 | 0.0024 |
| PERM | Myeloperoxidase | 0.33 |
| IL18 | Interleukin-18 | 0.2 |
| ELNE | Neutrophil elastase | 0.29 |
| BPIB1 | BPI fold-containing family B member 1 | 0.15 |
| FHR1 | Complement factor H-related protein 1 | 0.4 |
| RNAS2 | Non-secretory ribonuclease | 0.27 |
| LYSC | Lysozyme C | 0.65 |
| CFAB | Complement factor B | 0.52 |
| CATG | Cathepsin G | 0.96 |
| ATRN | Attractin | 0.18 |
| PTMA | Prothymosin alpha | 0.79 |
| NGAL | Neutrophil gelatinase-associated lipocalin | 0.84 |
| VTNC | Vitronectin | 0.64 |
| ICAM1 | Intercellular adhesion molecule 1 | 0.0023 |
| IL1AP | Interleukin-1 receptor accessory protein | 0.0032 |
| CDC42 | Cell division control protein 42 | 0.0022 |
| IL6RB | Interleukin-6 receptor subunit beta | 0.0051 |
| PIGR | Polymeric immunoglobulin receptor | 0.00038 |
| CO4B | Complement C4-B | 0.028 |

#### Anti-Oxidation Proteins

| Gene ID | Name | t test |
|---|---|---|
| CATA | Catalase | < 0.0001 |
| GSHR | Glutathione reductase, mitochondrial | 0.00057 |
| NQO1 | NAD(P)H dehydrogenase [quinone] 1 | 0.093 |
| SODC | Superoxide dismutase [Cu-Zn] | 0.11 |
| AK1A1 | Alcohol dehydrogenase [NADP(+)] | 0.12 |
| GPX1 | Glutathione peroxidase 1 | 0.052 |
| GSTA1 | Glutathione S-transferase alpha 1 | 0.51 |
| GPX4 | Glutathione peroxidase 4 | 0.75 |
| CBR3 | Carbonyl reductase [NADPH] 3 | 0.12 |
| AL9A1 | 4-trimethylaminobutyraldehyde dehydrogenase | 0.0092 |
| ANXA1 | Annexin A1 | 0.33 |
| CBR1 | Carbonyl reductase [NADPH] 1 | 0.088 |
| ADHX | Alcohol dehydrogenase | 0.0043 |
| HSPB1 | Heat shock protein beta-1 | 0.0043 |
| LDHA | L-lactate dehydrogenase A | 0.0052 |
| PRDX6 | Peroxiredoxin-6 | 0.0011 |
| LDHB | L-lactate dehydrogenase B | < 0.0001 |
| AL7A1 | Alpha-aminoadipic semialdehyde dehydrogenase | 0.00047 |
| PRDX5 | Peroxiredoxin-5 | 0.00065 |
| ARK72 | Aflatoxin B1 aldehyde reductase member 2 | < 0.0001 |

### Table S2

#### Antimicrobial Proteins

| Gene ID | Name | t test |
|---|---|---|
| MUC5A | Mucin-5AC | < 0.0001 |
| S10A8 | Protein S100-A8 | 0.29 |
| S10A9 | Protein S100-A9 | 0.32 |
| S10A7 | Protein S100-A7 | 0.51 |
| MUC4 | Mucin-4 | 0.13 |
| LYSC | Lysozyme C | 0.65 |
| DEF1 | Neutrophil defensin 1 | 0.63 |
| MUC1 | Mucin-1 | 0.85 |
| SFPA2 | Pulmonary surfactant-associated protein A2 | 0.12 |
| CAMP | Cathelicidin antimicrobial peptide | 0.44 |
| PGRP2 | N-acetylmuramoyl-L-alanine amidase | 0.14 |
| CO6 | Complement component C6 | 0.092 |
| SFTPD | Pulmonary surfactant-associated protein D | 0.0081 |
| CO4A | Complement C4-A | 0.0061 |
| CO2 | Complement C2 | < 0.0001 |
| CO5 | Complement C5 | 0.011 |
| CO3 | Complement C3 | 0.00034 |

#### Complement and Surfactant Proteins

| Gene ID | Name | t test |
|---|---|---|
| FHR1 | Complement factor H-related protein 1 | 0.4 |
| CFAH | Complement factor H | 0.13 |
| CFAB | Complement factor B | 0.52 |
| CO8G | Complement component C8 | 0.95 |
| SFPA2 | Pulmonary surfactant-associated protein A2 | 0.12 |
| C1R | Complement C1r | 0.72 |
| C1S | Complement C1s | 1 |
| CFAI | Complement factor I | 0.23 |
| CO9 | Complement component C9 | 0.12 |
| C4BPA | C4b-binding protein alpha chain | 0.66 |
| C1RL | Complement C1r subcomponent-like protein | 0.048 |
| CO6 | Complement component C6 | 0.092 |
| CO7 | Complement component C7 | 0.16 |
| CO8B | Complement component C8 | 0.028 |
| SFTPD | Pulmonary surfactant-associated protein D | 0.0081 |
| CO4A | Complement C4-A | 0.0061 |
| CFAD | Complement factor D | 0.0026 |
| CO2 | Complement C2 | < 0.0001 |
| CO4B | Complement C4-B | 0.028 |
| C5 | Complement C5 | 0.011 |
| CO3 | Complement C3 | 0.00034 |
| C1QC | Complement C1q subcomponent subunit C | 0.0056 |
| C1QA | Complement C1q subcomponent subunit A | 0.0061 |
| C1QB | Complement C1q subcomponent subunit B | 0.0031 |

### Table S3

#### IG and IG Receptor Proteins

| Gene ID | Name | t test |
|---|---|---|
| IGHD | Isoform 2 of Immunoglobulin heavy constant delta | 0.28 |
| IGHM | Immunoglobulin heavy constant mu | 0.55 |
| IGHG3 | Immunoglobulin heavy constant gamma 3 | 0.36 |
| IGHG1 | Immunoglobulin heavy constant gamma 1 | 0.7 |
| IGKC | Immunoglobulin kappa constant | 0.42 |
| IGHA2 | Immunoglobulin heavy constant alpha 2 | 0.27 |
| IGHG2 | Immunoglobulin heavy constant gamma 2 | 1 |
| FCGBP | IgGFc-binding protein | 0.028 |
| IGHA1 | Immunoglobulin heavy constant alpha 1 | 0.016 |
| IGHG4 | Immunoglobulin heavy constant gamma 4 | 0.096 |
| IGJ | Immunoglobulin J chain | 0.0066 |
| PIGR | Polymeric immunoglobulin receptor | 0.00038 |

#### Hydrolytic Proteins

| Gene ID | Name | t test |
|---|---|---|
| BGAL | Beta-galactosidase | 0.00088 |
| PPA5 | Tartrate-resistant acid phosphatase type 5 | 0.00019 |
| LYAG | Lysosomal alpha-glucosidase | < 0.0001 |
| PERM | Myeloperoxidase | 0.33 |
| GPX3 | Glutathione peroxidase 3 | 0.28 |
| GPX1 | Glutathione peroxidase 1 | 0.052 |
| GPX4 | Glutathione peroxidase 3 | 0.75 |
| PPBT | Alkaline phosphatase | < 0.0001 |
| IAH1 | Isoamyl acetate-hydrolyzing esterase 1 | < 0.0001 |
| GANAB | Neutral alpha-glucosidase AB | 0.02 |
| MA1A1 | Mannosyl-oligosaccharide 1,2-alpha-mannosidase IA | 0.00054 |

**Figure S1**

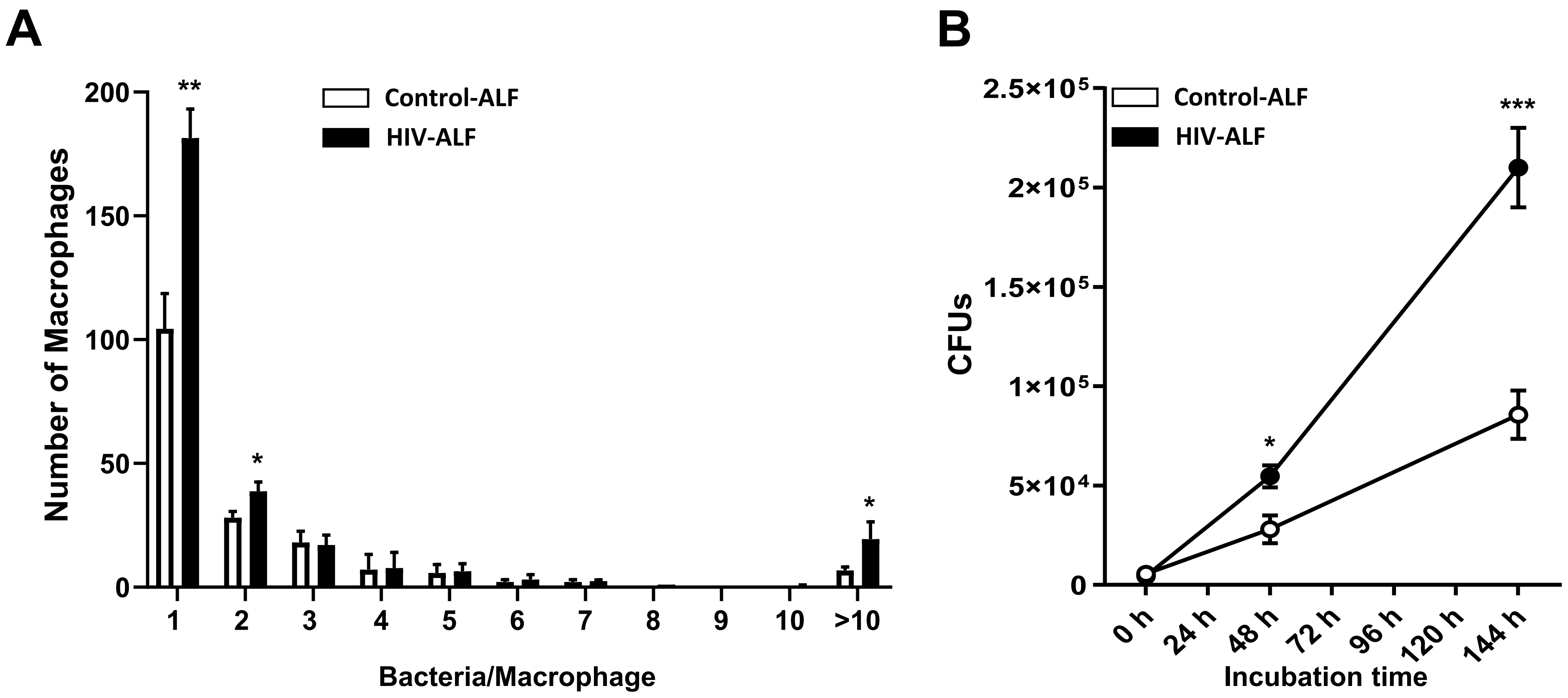
A
Number of Macrophages
200
150
100
50
0
**
*
*
Control-ALF
HIV-ALF
1
2
3
4
5
6
7
8
9
10
>10
Bacteria/Macrophage
B
CFUs
2.5×10⁵
2×10⁵
1.5×10⁵
1×10⁵
5×10⁴
0
Control-ALF
HIV-ALF
***
*
0 h
24 h
48 h
72 h
96 h
120 h
144 h
Incubation time
